# Urgent Brain Vascular Regeneration Occurs via Lymphatic Transdifferentiation

**DOI:** 10.1101/2021.04.23.441119

**Authors:** Jingying Chen, Xiuhua Li, Rui Ni, Qi Chen, Qifen Yang, Jianbo He, Lingfei Luo

## Abstract

Acute ischemic stroke damages regional brain blood vessel (BV) network. Urgent recovery of basic blood flows, which represents the earliest regenerated BVs, are critical to improve clinical outcomes and minimize lethality. Although the late-regenerated BVs have been implicated to form via growing along the meninge-derived lymphatic vessels (iLVs), mechanisms underlying the early, urgent BV regeneration remain elusive. Using zebrafish cerebrovascular injury models, we show that the earliest regenerated BVs come from lymphatic transdifferentiation, a hitherto unappreciated process in vertebrates. Mechanistically, LV-to-BV transdifferentiation occurs exclusively in the stand-alone iLVs through Notch activation. In the track iLVs adhered by nascent BVs, transdifferentiation never occurs because the BV-expressing EphrineB2a paracellularly activates the iLV-expressing EphB4a to inhibit Notch activation. Suppression of LV-to-BV transdifferentiation blocks early BV regeneration and becomes lethal. These results demonstrate that urgent BV regeneration occurs via lymphatic transdifferentiation, suggesting this process and key regulatory molecules EphrinB2a/EphB4a/Notch as new post-ischemic therapeutic targets.

## INTRODUCTION

Acute ischemic stroke, one of the leading causes of mortality and adult disability worldwide, develops when BVs supplying oxygen and nutrients to certain brain areas are occluded (Prabhakaran et al., 2015). It results in damages to regional vascular network and brain tissues. Reduction of brain injury together with promotion of neoangiogenesis and neurogenesis appear to be the most promising therapeutic strategy (Hermann and Chopp, 2012). According to this strategy, resolvation of the stroke-caused edema and prompt recovery of basic blood flows to supply minimal oxygen and nutrients serve as two of the major principles to alleviate symptoms and minimize lethality (Krupinski et al., 1993; Krupinski et al., 1994; Hayashi et al., 2003; Bardutzky and Schwab, 2007; Arai et al., 2009; Ergul et al., 2012).

Roles of lymphatic vessels (LVs) in anti-edema and the late wave of BV regeneration after cerebrovascular damages have lately been highlighted (Chen et al., 2019; Yanev et al., 2020). Traditionally, the lymphatic system is responsible for the maintenance of fluid homeostasis, fat absorption, and immune surveillance (Alitalo et al., 2005). During development, the lymphatic endothelial cells (LECs) are derived from blood vessel endothelial cells (BECs) in vertebrates (Sabin, 1902; Yaniv et al., 2006; Srinivasan et al., 2007; Schulte-Merker et al., 2011; Nicenboim et al., 2015). The lymphatics inside the skull, including mammalian meningeal lymphatic vessels (LVs) and leptomeningeal LECs (LLECs), as well as zebrafish brain LECs (BLECs) also called meningeal mural LECs (muLECs) or fluorescent granule perithelial cells (FGPs), have recently been discovered (Louveau et al., 2015; Aspelund et al., 2015; Antila et al., 2017; Bower et al., 2017; van Lessen et al., 2017; Venero Galanternik et al., 2017; Shibata-Germanos et al., 2020). In response to brain vascular injury, BLECs rapidly ingrown into the injured parenchyma and form lumenized iLVs, which on one hand drain interstitial fluid to resolve brain edema, on the other hand act as “growing tracks” of nascent BVs (Chen et al., 2019). However, the iLV-guided growth of nascent BVs is a time-consuming process, which serves as the mechanism of the late wave of BV regeneration for the rebuilt of cerebrovascular network, but cannot meet the urgent requirements of basic blood flows in the injured brain. Cellular and molecular mechanisms underlying prompt formation of early-regenerated BVs still remain mystery.

In the present study, we applied genetic ablation and photochemical thrombosis models to induce injuries to brain vasculature in zebrafish larvae and adults. We found that the earliest regenerated BVs arise from iLV transdifferentiation. The iLVs can be subdivided into track iLVs and stand-alone iLVs based on acting as and not acting as the “growing tracks” of nascent BVs, respectively. The LV-to-BV transdifferentiation occurs exclusively in the stand-alone iLVs through activation of Notch signaling, but never occurs in the track iLVs because the BV-expressing EphrineB2a paracellularly activates the iLV-expressing EphB4a to inhibit Notch activation. Suppression of the stand-alone iLV-to-BV transdifferentiation blocks the formation of early-regenerated BVs and becomes lethal. This study reveals a previously unexpected LV-to-BV transdifferentiation process in vertebrate, which occurs under pathological circumstances and sheds lights on how the urgent recovery of post-ischemic basic blood flows is achieved.

## RESULTS

### Early-Regenerated BVs Are Converted from Stand-Alone iLVs

The brain parenchyma is physiologically devoid of LVs. We have previously reported that only a portion of iLVs, but not all of them, undergo apoptosis once rebuilt of brain vasculature is approaching (Chen et al., 2019). This phenomenon makes us hypothesize the contribution of non-apoptotic iLVs to the early-regenerated BVs for the prompt recovery of basic blood flows. Before exploration of this hypothesis, we first analyzed the specificity of *lyve1b* promoter inside the skull under physiological and cerebrovascular injury circumstances, in addition to previous reports showing that brain and meningeal blood vessels are negative for *lyve1b* (Bower et al., 2017; Chen et al., 2019; van Lessen et al., 2017; Venero Galanternik et al., 2017). Although *lyve1b* has previously been reported to express in both LECs and some trunk veins (Okuda et al., 2012), all the *lyve1b:DsRed*-expressing BLECs and iLVs inside the skull were positive for anti-Prox1 in both control and Mtz-treated larvae at 2 days post treatment (dpt), equivalent to 5 days post fertilization (dpf) (Figures S1A, S1B, and S1D). Prox1 is a marker of LEC, but not BEC (Koltowska et al., 2015; Gancz et al., 2019). None of these anti-Prox1+*lyve1b:DsRed*+ cells expressed the BEC-specific *kdrl:Dendra2* transgene at 2 dpt (Figures S1A and S1B). Additionally, at 2 dpt after Mtz treatment, the *lyve1b:GFP*-expressing vessels in the brain completely overlapped with the *prox1a:KalTA4;UAS:TagRFP*-expressing vessels (van Impel et al., 2014) (Figures S1C and S1D). Furthermore, after incubation of 4-hydroxytamoxifen (4-OHT) with the *Tg(kdrl:DenNTR; lyve1b:CreER^T2^; β-actin2:loxP-STOP-loxP-DsRed)* lineage tracing line from 3 dpf to 4 dpf, all the brain BVs were negative for DsRed at 7 dpf and 9 month post fertilization (mpf) (Figures S1E-S1G). All these evidences indicate that *lyve1b* is entirely specific to LECs and not expressed in any BEC in the meninx and brain, suggesting *lyve1b* as a reliable LEC-specific marker inside the skull.

To explore the fate of non-apoptotic iLVs, we continue using the *Tg(kdrl:DenNTR; lyve1b:CreER^T2^; β-actin2:loxP-STOP-loxP-GFP)* triple transgenic line to perform lineage tracing. This line was crossed with *Tg(lyve1b:DsRed)*, followed by treatment with Mtz as the negative control and treatment with DMSO plus 4-OHT as the positive control (Figures S1H-S1J), ensuring no Mtz-caused CreER leakiness and labeling efficiency of the LEC-derived cells, respectively. When the triple transgenic larvae were incubated with Mtz plus 4-OHT and analyzed at 4 dpt, 28.8±3.0% of the GFP-positive, LEC-derived vessels were also positive for the BEC-specific Dendra2 (Figures 1A-1D).

**Figure 1.**
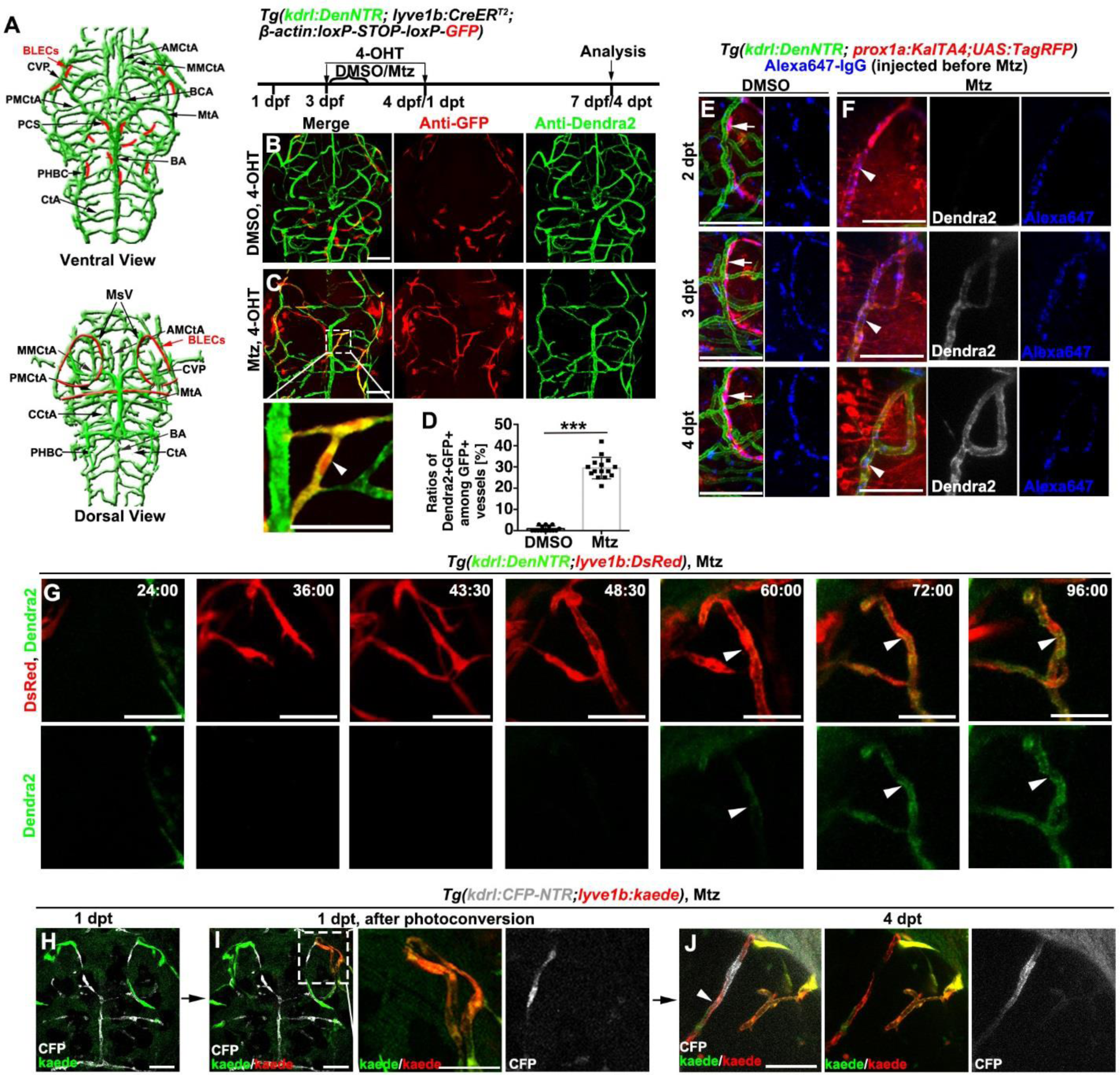
Early brain BV regeneration occurs via iLV-to-BV transdifferentiation. (A) The ventral and dorsal views of zebrafish brain vasculature at larval stages, which include all the imaging areas of high magnification pictures. Green and red represent brain BVs and BLECs, respectively. (B-D) Controlled by uninjured larvae (B), a portion of GFP+ vessels were positive for Dendra2 after Mtz treatment (C), which is shown in higher magnification in the framed area (arrowhead). (D) The statistics show ratios of the Dendra2+GFP+ vessels among GFP+ vessels (n=15 larvae, Two-tailed unpaired *t*-test, ***, *p*<0.0001). (E and F) In contrast to the RFP+Alexa647+ BLECs in the uninjured control (E, arrows), the RFP+Alexa647+ iLVs gradually expressed Dendra2 from 2 dpt to 4 dpt after Mtz treatment (F, arrowheads). The Alexa647-IgG was injected before Mtz treatment. See also Video S1. (G) Time-lapse imaging showed the real-time processes of iLV ingrowth and gain of Dendra2 fluorescence. The expression of Dendra2 became detectable from 60:00/60 hpt. Arrowheads indicate the transdifferentiating iLVs. The elapsed time is indicated in hours:minutes after 0 dpt. See also Video S2. (H-J) BLECs were labeled by the Kaede-green at 1 dpt shortly before ingrowth (H). The photoconverted BLECs were marked by the white frame in (I) and enlarged in right panels. At 4 dpt, one iLV derived from the Kaede-red BLECs expressed the BEC-specific CFP (J, arrowhead). See also Video S3. Scale bar, 50 μm. Data are represented as mean±SD. See also Figures S1 and S2. AMCtA, anterior (rostral) mesencephalic central artery; PCS, posterior (caudal) communicating segment; BCA, basal communicating artery; BA, basilar artery; CCtA, cerebellar central artery; CtA, central artery; CVP, choroidal vascular plexus; MMCtA, middle mesencephalic central artery; MsV, mesencephalic vein; MtA, metencephalic artery; PHBC, primordial hindbrain channel; PMCtA, posterior (caudal) mesencephalic central artery; BLECs, brain lymphatic endothelial cells.

These data first suggest that a portion of iLVs could be converted to BVs. Then, another lineage tracing line *Tg(kdrl:DenNTR; lyve1b:CreER^T2^; kdrl:loxP-STOP-loxP-H2b-GFP)* specific for the detection of the LEC-derived BECs, was applied. At 4 dpt after injury, 32.9±6.3% of brain BVs turned out to be positive for H2b-GFP (Figures S2A-S2D), substantiating that a portion of regenerating BVs originated from iLVs.

Besides the lineage tracing assays above, we also carried out three independent live imaging experiments to validate the iLV-to-BV transdifferentiation. First, the BLECs and ingrown LECs (iLECs), but not BECs, are able to endocytose and accumulate the Alexa647-IgG macromolecules (Chen et al., 2019; van Lessen et al., 2017). The *lyve1b:DsRed* or *prox1a:KalTA4;UAS:TagRFP* transgene, together with endocytosed Alexa647-IgG, were applied to label LECs for live imaging analyses (Figures 1E and S2E). Live imaging showed that the expression of BEC-specific Dendra2 was absent in the iLVs at 2 dpt, but became present at 3 dpt. The Alexa647-IgG injected before Mtz treatment continuously maintained in the transdifferentiating endothelial cells from 2 dpt to 4 dpt (Figures 1F; Figures S2F). The iLV-derived BVs obtained blood flows at 3 dpt (Video S1; Figure S2F, arrowhead), which represented the earliest formation of functional BVs and the promptest recovery of blood flows in the injured brain parenchyma. Second, time-lapse imaging using the *Tg(kdrl:DenNTR; lyve1b:DsRed)* line showed the real-time processes of single *lyve1b*+ iLV ingrowth and gain of the *kdrl:Dendra2* fluorescence (Figure 1G; Video S2). Ingrowth of BLECs to form iLVs occurred from 24 hours post Mtz treatment (hpt)/1 dpt. The ingrown iLVs began to express the BEC-specific Dendra2 at 60 hpt/2.5 dpt, which became evident at 72 hpt/3 dpt, displaying the real-time process of iLV-to-BV transdifferentiation. Third, under the photoconvertible *Tg(kdrl:CFP-NTR; lyve1b:kaede)* transgenic background, the green-to-red fluorescence photoconversion was applied to a small portion of BLECs shortly before ingrowth at 1 dpt. At 4 dpt, one of the iLVs derived from the Kaede-red BLECs expressed the BEC-specific CFP and obtained blood flows (Figures 1H-1J; Video S3), indicating the origination of this BV from the photoconverted BLECs. All these results demonstrate that a portion of iLVs are directly converted to functional BVs, achieving the earliest formation of post-injured functional brain BVs.

We then investigate whether the vessels undergoing iLV-to-BV conversion switch their molecular and structural identities accordingly. At 3 dpt, the transdifferenating iLVs double positive for lyve1b:DsRed and kdrl:Dendra2 lost the lymphatic markers anti-Prox1 and *vegfr3* (Bower et al., 2017; van Lessen et al., 2017; Venero Galanternik et al., 2017) (Figures 2A-2C). The lineage tracing system showed that the iLV-derived BVs obtained the blood-brain barrier marker *glut1b* (Zheng et al., 2010) (Figures 2D and 2E). The correlative light and electron microscopy (CLEM) and focused ion beam-scanning electron microscopy (FIB-SEM) showed that the DsRed+Dendra2+ vessels at 4 dpt and the DsRed-Dendra2+ BVs at 6 dpt owned similar mural structures and blood cells in the lumen. By contrast, the DsRed+Dendra2-iLVs at 2 dpt exhibited lymphatic structures, much thinner mural and no blood cell content (Figures 2F-2H). Live imaging in the *Tg(kdrl:DenNTR; lyve1b:DsRed; pdgfrb:GFP)* transgenic larvae indicated that the BV and DsRed+Dendra2+ vessel at 4 dpt, but not its DsRed+Dendra2-state prior to transdifferentiation at 2 dpt, recruited the GFP+ pericytes (Ando et al., 2016) (Figures 2I and 2J, arrowheads). These results demonstrate that the transdifferentiating iLVs switch their molecular and structural identities from LV’s to BV’s.

**Figure 2.**
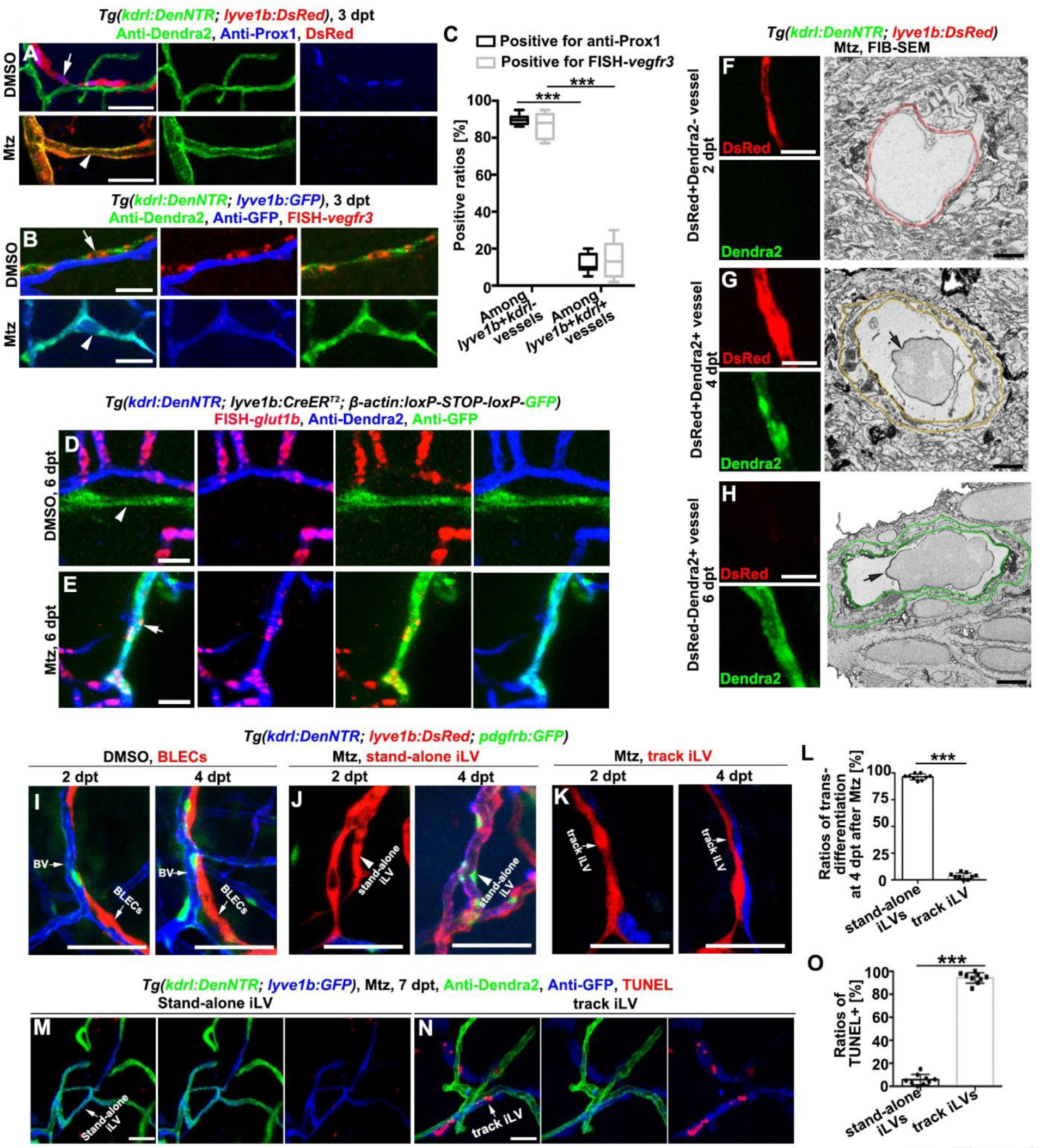
The transdifferenting iLVs switch molecular and structural identities and are restricted to stand-alone iLVs. (A–C) The *lyve1b*+*kdrl*-BLECs in the uninjured control (arrows), but not the transdifferentiating *lyve1b*+*kdrl*+ iLVs at 3 dpt after injury (arrowheads), were positive for anti-Prox1 (A) and FISH-*vegfr3* (B). The statistics show the ratios of vessels positive for anti-Prox1 or FISH-*vegfr3* among all the *lyve1b*+*kdrl*-vessels and all the *lyve1b*+*kdrl*+ vessels. (C, n=6 larvae. Two-way ANOVA by Dunnett’s multiple comparisons test. ***, *P*<0.0001). Scale bar, 20 μm. (D and E) Positively controlled by brain BVs in the uninjured larvae (D), The lineage tracing system indicated that the vessels double positive for GFP and Dendra2 (E, arrow) and single positive for Dendra2 (D and E), but not vessels single positive for GFP (D, arrowhead), expressed the blood brain barrier marker *glut1b*. Scale bar, 20 μm. (F-H) Single FIB-SEM image planes (right row) indicate cross sections of the vessels shown in the left row. Note that the mural of the DsRed+Dendra2+ vessel (G) is similar to the DsRed-Dendra2+ BV (H), but much thicker than the DsRed+Dendra2-iLV (F). Color rings mark the inner and outer surfaces of murals. Arrows indicate blood cells. Scale bar, 20 μm and 1 μm. (I-L) Live imaging shows BV, BLECs (I), stand-alone iLVs (J, arrowheads), and track iLVs (K) at 2 dpt and 4 dpt. Note that similar with the BV (I), only the stand-alone iLVs express Dendra2 and recruit the GFP+ pericytes at 4 dpt after Mtz (J, arrowheads). (L) The statistics show the ratios of transdifferentiation in the stand-alone iLVs and track iLVs at 4 dpt (n=9 larvae, Two-tailed unpaired *t*-test, ***, *p* <0.0001). Scale bar, 50 μm. (M-O) The TUNEL signals in the stand-alone iLVs (M) and track iLVs (N) at 7 dpt and their statistics (O, n=9 larvae, Two-tailed unpaired *t*-test, ***, *p*<0.0001). Scale bar, 50 μm. Data are represented as mean±SD. See also Figure S2.

The pericyte-labeled line and live imaging studies above hypothesize occurrence of transdifferentiation exclusively in the stand-alone iLVs, which meant no adhersion of nascent BVs. So, we examined a large number of individual iLVs to identify the subpopulations destined to transdifferentiation. Only the stand-alone iLVs were able to be converted to early-formed, pericyte-covered BVs (Figures 2J and 2M). And vice versa, iLVs acting as “growing tracks” of nascent BVs, hereafter called track iLVs, never recruited pericytes and finally underwent apoptosis (Figures 2K, 2L, 2N, and 2O).

Therefore, iLVs could be subdivided into two populations, track iLVs and stand-alone iLVs, which finally underwent apoptosis and transdifferentiation, respectively.

Comparing to the stand alone iLV-converted BVs that became functional within 2 days after injury (1 dpt to 3 dpt), nascent BVs growing along the track iLVs represented the late wave of BV regeneration, which took approximately 7 days (1 dpt to 8 dpt) (Chen et al., 2019).

### EphB4a Paracellularly Activated by EphrinB2a Inhibits iLV Transdifferentiation

To enable manipulations of the iLV-to-BV transdifferentiation process to understand the biological significance of early-regenerated BVs, we first need to decipher the underpinning mechanisms. The data above have hypothesized that the adhering nascent BVs might provide major inhibitory factors to block the transdifferentiation of track iLVs. The most reasonable molecules that could afford the BEC-iLEC paracellular inhibitory effects appeared to be the paracellular membrane ligand/receptor. So, we screened these pairs of ligands/receptors for their respective expressions in nascent BVs and track iLVs, among which EphrinB2a/EphB4a were identified (Adams et al., 1999). At 3 dpt, the receptor *ephB4a* and the ligand *ephrinB2a* were specifically expressed in the track iLVs and the adhering nascent BVs, respectively, while they were absent in the uninjured brain (Figures 3A-3D). The detection of tyrosine-phosphorylated EphB4 indicated the activation of EphB4 in the track iLVs, but not in the stand-alone iLVs (Figure 3E-3G).

**Figure 3.**
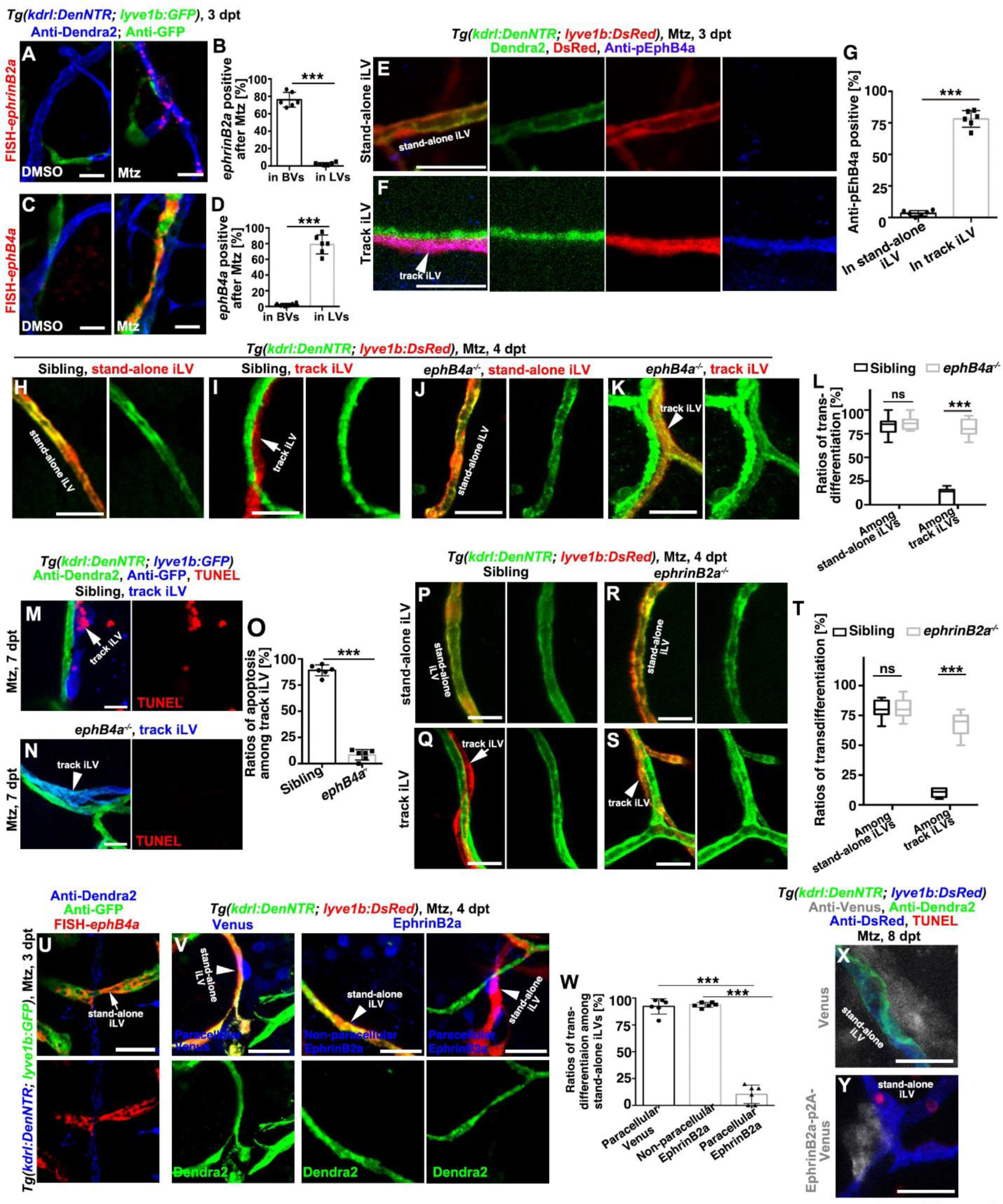
EphrinB2a/EphB4a are necessary and sufficient to inhibit the iLV-to-BV transdifferentiation. (A-D) In contrast to their absence in the brain of uninjured control, *ephrinB2a* and *ephB4a* was respectively activated in the Dendra2+GFP-BVs and Dendra2-GFP+ LVs after Mtz treatment (A and C). The statistics show *ephrinB2a* and *ephB4a* expression in the vessels, respectively (B and D, n=6 larvae, Two-tailed unpaired *t*-test, ***, *p*<0.0001). (E-G) In contrast to the stand-alone iLV (E), expression of the tyrosine phosphorylated EphB4 (pEphB4) indicate the activation of EphB4a in the track iLVs of the injured brain (F). The statistics show the ratios of pEphB4-positive among stand-alone and track iLVs (G, n=6 larvae, Two-tailed unpaired *t*-test. ***, *p*<0.0001). (H-L) After Mtz treatment, the stand-alone iLV (H), but not the track iLVs (I, arrow), underwent transdifferentiation in siblings. In the *ephB4a* mutant, both stand-alone (J) and track iLVs expressed Dendra2 (K, arrowhead). The statistics show the ratios of transdifferentiation among stand-alone iLVs and track-iLVs in siblings and in the *ephB4a* mutant. (L, n=7 larvae, Two-way ANOVA by Sidak’s multiple comparisons test. ns, *P*=0.7299; ***, *P*<0.0001). See also Video S3. (M-O) After Mtz treatment, the track iLVs finally underwent apoptosis (M, arrow). In the *ephB4a* mutant, the track iLVs expressed Dendra2 and became TUNEL-negative (N, arrowhead). The statistics show the ratios of apoptosis among the track iLVs in sibling and in the *ephb4a* mutant (O, n=6 larvae, Two-tailed unpaired *t*-test, ***, *p*<0.0001). (P-T) In contrast to siblings (Q, arrow), the track iLVs in the *ephrinB2a* mutant expressed Dendra2 (S, arrowhead), whereas the transdifferentiation of stand-alone iLVs remained unaffected (P and R). The statistics show the ratios of transdifferentiation among stand-alone iLVs and track-iLVs in sibling and in the *ephrinB2a* mutant (T, n=7 larvae. Two-way ANOVA by Sidak’s multiple comparisons test. ns, *p*=0.9675; ***, *p*<0.0001). (U-W) The *ephB4a* expressed in the stand-alone iLVs after Mtz treatment (U). The paracellularly expressed EphrinB2a, but not the non-paracellular EphrinB2a or paracellular Venus, repressed the Dendra2 expression in the stand-alone iLVs (V). The statistics show the ratios of transdifferentiation in the subpopulations of stand-alone iLVs (W, n=6 larvae, Two-tailed unpaired *t*-test, ***, *p*<0.0001). (X andY) The paracellularly expressed EphrinB2a (Y), but not Venus (X), caused apoptosis of the stand-alone iLVs. Scale bar, 20 μm. NS, not significant. Data are represented as mean±SD. See also Figures S3, S4, S5, and S7.

Roles of EphrinB2a/EphB4a in the iLV transdifferentiation were analyzed using the *ephB4a^hu3445^* mutant (Choe and Crump, 2015). The mutant exhibited normal vascular and lymphatic development (Figures S3A and S3B). After brain vascular injury, the track iLVs never transdifferentiated at 4 dpt and underwent apoptosis at 7 pdt in the control siblings (Figures 3I and 3M). In the *ephB4a* mutant, the natural transdifferentiation of stand-alone iLVs was unaffected (Figures 3H and 3J). However, the track iLVs in the mutant also exhibited transdifferentiation and failed to undergo apoptosis, and were finally converted to BVs with blood flows (Figures 3K, 3L, 3N, 3O; Video S4). The transdifferentiation of track iLVs became observable from 4 dpt (Figure S4). The inhibitory effect of EphB4a on the transdifferentiation of iLVs was validated by the construction of a constitutively active form EphB4aEE with double mutations of Tyr-596 and Tyr-602 into glutamate (Zisch et al., 2000; Yang et al., 2006). The heat shock-induced, LEC-specific overexpression of EphB4aEE blocked the transdifferentiation of stand-alone iLV (Figure S5). These results demonstrate that the track iLEC-expressing EphB4a is active to inhibit the track iLV-to-BV transdifferentiation.

We next investigate whether the inhibitory activity of EphB4a on the track iLV transdifferentiation is activated by EphrinB2a. An *ephrinB2a* mutant, in which frame shift and protein truncation occurred at amino acid 6 and 49, respectively (Figure S3C), was generated. Cerebrovascular and BLECs development remained normal in the *ephrinB2a* mutant (Figures S3D and S3E). Ectopic transdifferentiation of track iLVs was observed in the *ephrinB2a* mutant, without affecting the natural transdifferentiation of stand-alone iLVs (Figures 3P-3T). These results indicate the repressive role of EphrinB2a/EphB4a on the track iLV transdifferentiation, so that the defective EphB4a or EphrinB2a leads to derepressed transdifferentiation of track iLVs.

After brain vascular injury, the expression of *ephB4a* was not only present in the track iLVs (Figure 3C), but also in the stand-alone iLVs (Figure 3U). To explore whether the paracellular EphrinB2a/EphB4a is sufficient to inhibit the iLV-to-BV conversion, the *hsp70l:ephrinB2a-p2A-Venus* plasmid or the *hsp70l:Venus* control plasmid was injected and heat-shocked to generate mosaic larvae. The paracellularly expressed EphrinB2a, but not the non-paracellular EphrinB2a or paracellular Venus, blocked the transdifferentiation of stand-alone iLVs (Figures 3V and 3W). These untransdifferentiated stand-alone iLVs finally underwent apoptosis (Figures 3X and 3Y). All these data above demonstrate that EphrinB2a expressed by the nascent BECs paracellularly activates its receptor EphB4a on the track iLECs, which is necessary and sufficient to suppress the iLV-to-BV transdifferentiation.

### EphB4a Inhibits The iLV-to-BV Conversion via Blocking Notch

Low levels of Notch are required for the induction of lymphatic fate in venous BECs during development (Murtomaki et al., 2013), and Notch overexpression in the *in vitro* cultured LECs can reprogram the lymphatic to arterial cell fate (Kang et al., 2010). So, we investigated whether the repression of iLV-to-BV conversion by EphrinB2a/EphB4a is mediated by the suppression of Notch signaling. At 3 dpt after Mtz, expressions of the Notch ligan*d dll4*, the receptor *notch1a*, the downstream target gene *hey1*, and the functional Notch reporter *tp1:Venus* (Ninov et al., 2012) were activated in the stand-alone, transdifferentiating iLVs (Figures 4A, 4E, 4I, and 4M) and nascent BVs, but not in the track iLVs (Figures 4B, 4F, 4J, and 4N). By contrast, in the *ephB4a* mutant, track iLVs also exhibited *dll4*, *notch1a*, *hey1*, and *tp1:Venus* expressions (Figures 4C, 4D, 4G, 4H, 4K, 4L, 4O, and 4P). These data suggest that defective EphB4a leads to the derepression of Notch activities in the track iLVs.

**Figure 4.**
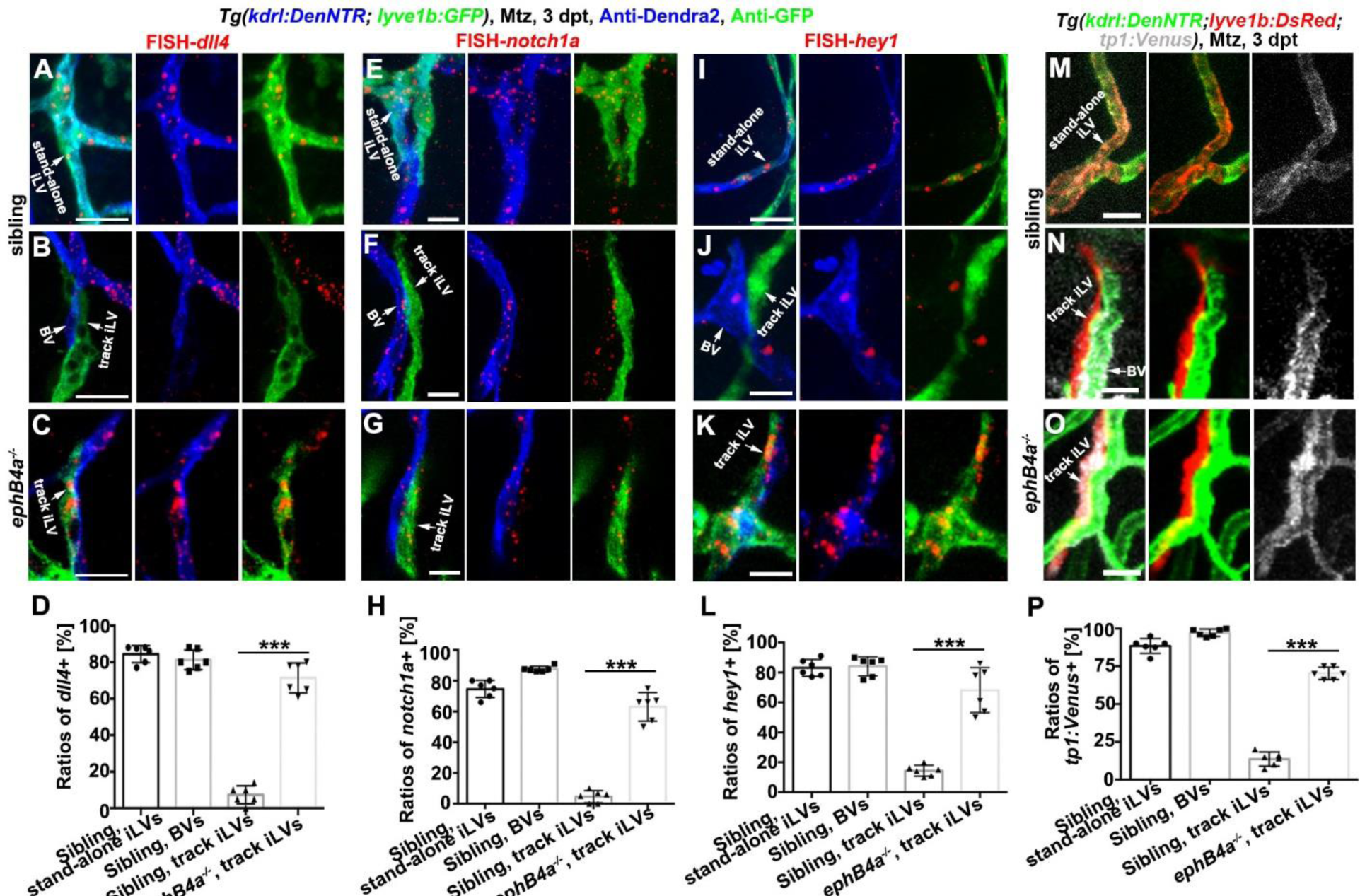
Defective EphB4a leads to derepression of Notch in the track iLVs. (A-L) Triple labeling of anti-Dendra2, anti-GFP, and FISH-*dll4* (A-C)/*notch1a* (E-G)/*hey1* (I-K). In the siblings, *dll4*/*notch1a*/*hey1* were activated in the Dendra2+GFP+ stand-alone iLVs (A, E, I) and Dendra2+GFP-nascent BVs (B, F, J), but not in the Dendra2-GFP+ track iLVs (B, F, J). By contrast, in the *ephB4a* mutant, the Dendra2-GFP+ track iLVs at 3 dpt also exhibited *dll4*, *notch1a*, and *hey1* expressions (C, G, K). The statistics show the *dll4*/*notch1a*/*hey1* expression in the vessels of sibling and *ephB4a* mutant (D, H, L, n=6 larvae, Two-tailed unpaired *t*-test, ***, *p*<0.0001). (M-P) The Notch functional reporter *tp1:Venus* was activated in the Dendra2+DsRed+ stand-alone iLVs (M) and Dendra2+DsRed-nascent BVs (N), but not in the Dendra2-DsRed+ track iLVs (N). By contrast, in the *ephB4a* mutant, the Dendra2-DsRed+ track iLVs at 3 dpt also exhibited Venus expression (O). The statistics show the *tp1:Venus* expression in the vessels of sibling and *ephB4a* mutant (P, n=6 larvae, Two-tailed unpaired *t*-test, ***, *p*<0.0001). Scale bar, 20 μm.. Data are represented as mean±SD. See also Figures S6 and S7.

Then, we examine whether the derepressed transdifferentiation of track iLVs in the *ephB4a* mutant (Figure 3K) was caused by the derepression of Notch activities. A well-known γ-secretase inhibitor DAPT that inhibits Notch signaling was applied (Lawson et al., 2001). The transdifferentiated track iLVs in the *ephB4a* mutant were rescued by the treatment with DAPT (Figures 5A-5C). Furthermore, rescue of the *ephB4a* mutant by the LEC-specific inhibition of Notch was performed using a dominant-negative isoform of the murine mastermind-like (dnMAML) protein, which incorporates into the Notch transcriptional complex but lacks the activity to recruit essential cofactors, thereby blocking the Notch activities (Zhao et al., 2014). Under the *Tg(kdrl:DenNTR; lyve1b:CreER^T2^; β-actin2:loxP-STOP-loxP-GFP; hsp70l:loxP-STOP-loxP-dnMAML-Flag)* transgenic background, heat-shock induction of the dnMAML-Flag expression specifically in the LEC-derived cells rescued track iLV transdifferentiation in the *ephB4a* mutant (Figures 5D-5G). These data indicate that EphrinB2a/EphB4a inhibit the iLV-to-BV transdifferentiation through suppression of Notch.

**Figure 5.**
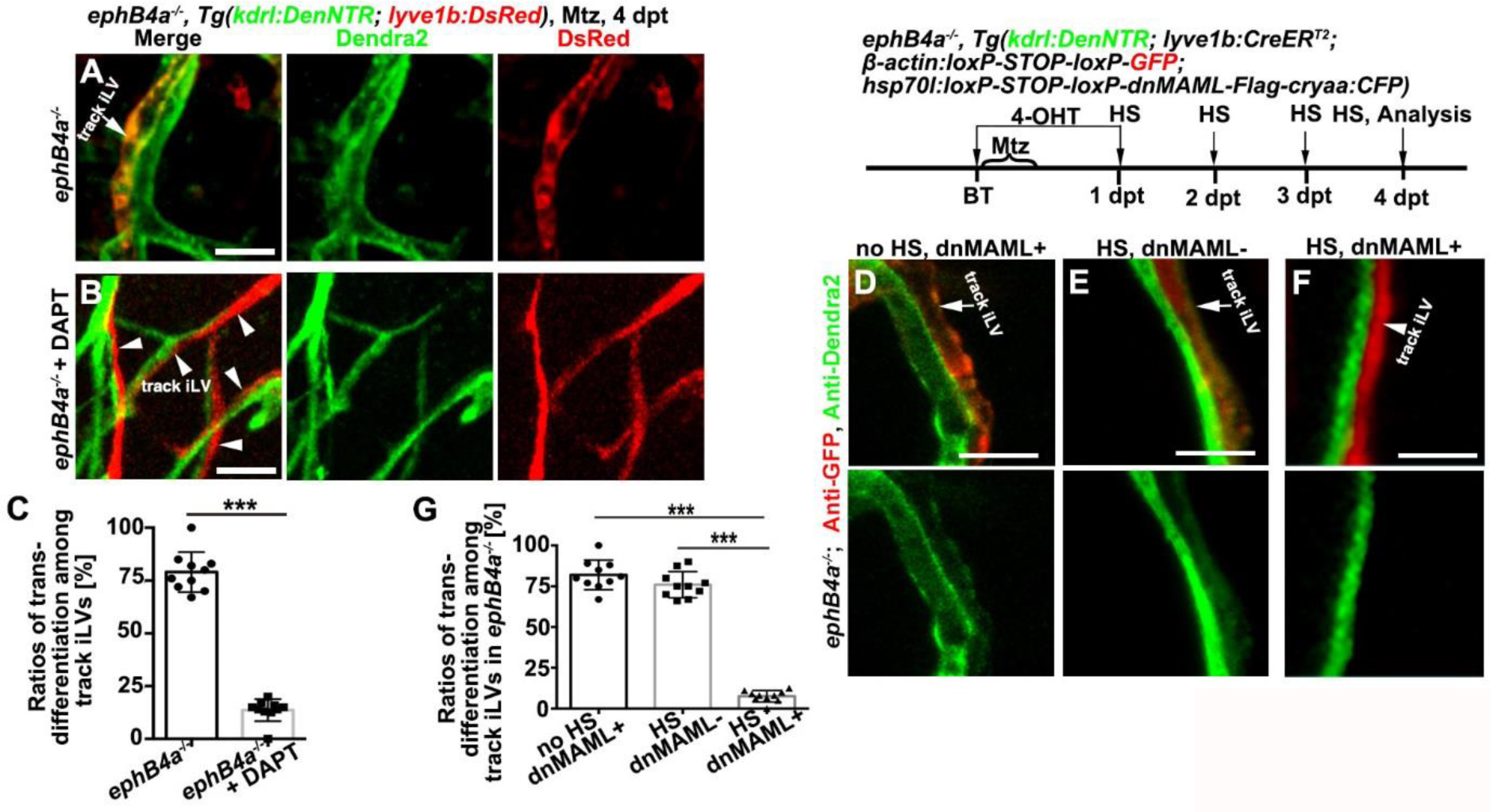
EphB4a represses track iLV transdifferentiation through suppression of Notch. (A-C) The derepressed transdifferentiation of track iLV (Dendra2+DsRed+) in the *ephB4a* mutant (A, arrow) was rescued by the treatment of DAPT, which returned to Dendra2-DsRed+ (B, arrowheads). The statistics show the ratios of transdifferentiation among track iLVs in the *ephB4a* mutant with or without DAPT treatment (C, n=10 larvae, Two-tailed unpaired *t*-test, ***, *p*<0.0001). (D-G) The derepressed transdifferentiation of track iLV (Dendra2+GFP+) in the *ephB4a* mutant without heat shock (D, arrow) or without dnMAML-Flag (E, arrow) was rescued by the heat shock-induced, LEC-specific overexpression of dnMAML-Flag, which exhibited Dendra2-GFP+ at 4 dpt (F, arrowhead). The statistics show the ratios of transdifferentiation among track iLVs in the *ephB4a* mutant groups (G, n=10 larvae, Two-tailed unpaired *t*-test, ***, *p*<0.0001). Scale bar, 20 μm. Data are represented as mean±SD. HS, heat shock. See also Figures S6 and S7.

### Early BV Regeneration Is Dependent on Notch and Enables Survival

The inhibition of track iLV-to-BV transdifferentiation by EphrinB2a/EphB4a via suppression of Notch together with activation of Notch in the stand-alone iLVs (Figures 4A, 4E, 4I, 4M) hypothesized that the transdifferentiation of stand-alone iLVs might be dependent on the Notch signaling. To exam this hypothesis, general inhibition of Notch signaling by DAPT or by a heat-shock-induced overexpression of dnMAML-GFP was applied. Either treatment significantly reduced the transdifferentiation of stand-alone iLVs (Figures S6A-S6F), which were validated and became even more evident when dnMAML-Flag was tissue-specifically overexpressed in the LEC-derived cells (Figures 6A-6C). And vice versa, general or LEC-specific activation of the Notch signaling using NICD-HA (Andersson et al., 2011) induced the track iLV-to-BV conversion (Figures S6G-S6I; Figures 6D-6F), indicating that Notch activation is sufficient to induce lymphatic transdifferentiation. All these data demonstrated that activation of Notch signaling in the iLECs is necessary and sufficient for the iLV-to-BV transdifferentiation.

**Figure 6.**
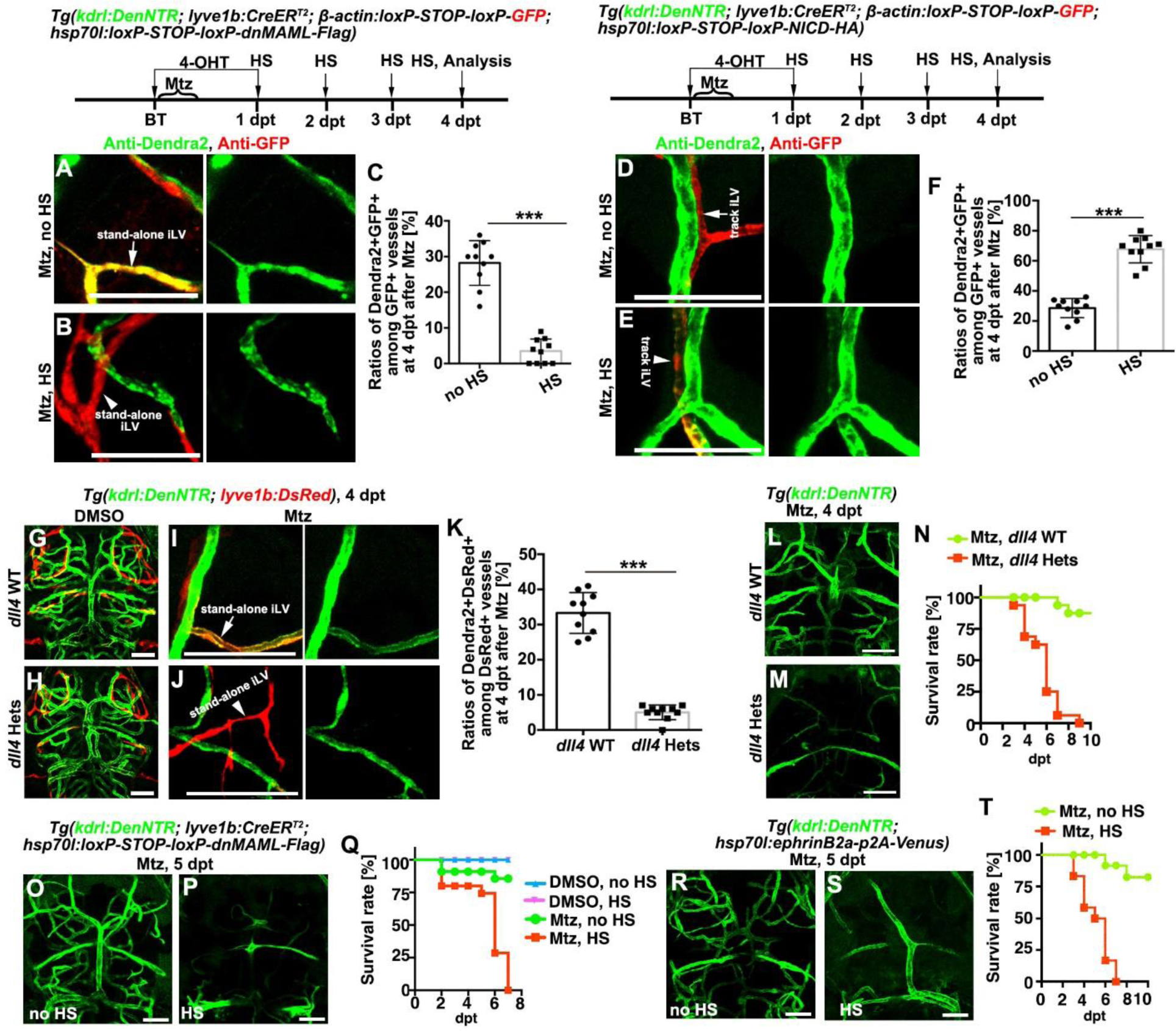
The stand-alone iLV-to-BV transdifferentiation is dependent on Notch and critical for post-injured survival. (A-C) Transdifferentiation of the stand-alone iLVs (A, arrow) was inhibited by the LEC-specific overexpression of dnMAML-Flag (B, arrowhead). The statistics show the ratios of Dendra2+GFP+ vessels among all the GFP+ vessels (C, n=10 larvae, Two-tailed unpaired *t*-test, ***, *p*<0.0001). (D-F) The track iLVs (D, arrows) were induced to undergo LV-to-BV transdifferentiation by the LEC-specific overexpression of NICD-HA (E, arrowhead). The statistics show the ratios of Dendra2+GFP+ vessels among all the GFP+ vessels (F, n=10 larvae, Two-tailed unpaired *t*-test, ***, *p*<0.0001). (G-K) In contrast to the wild-type (G), the uninjured *dll4* heterozygotes (H) were physiologically normal including brain BVs and BLECs. But after Mtz treatment, the stand-alone iLV-to-BV transdifferentiation (I, arrow) failed to occur in the *dll4* heterozygotes (J, arrowhead). The statistics show ratios of Dendra2+GFP+ vessels among all the GFP+ vessels (K, n=10 larvae, Two-tailed unpaired *t*-test, ***, *p*<0.0001). (L-T) The *dll4* heterozygotic mutation (L-N), or the LEC-specific overexpression of dnMAML-Flag (O-Q), or general overexpression of EphrinB2a (R-T), led to defects in the formation of early-regenerated BVs and became lethal. The statistics show the survival rates (N, n=32; Q, n=40; T, n=36). Note that the majority of the larvae with defective formation of early-regenerated BVs died before 6 dpt, while all of them died at 7 dpt. Scale bar, 50 μm.. Data are represented as mean±SD. Hets, heterozygotes. HS, heat-shock. See also Figures S6 and S7.

Finally, based on the mechanisms depicted above, we were able to manipulate the formation of early-regenerated BVs to assess their post-injured biological significance. Although the uninjured *dll4* heterozygous mutant was physiologically normal (Leslie et al., 2007) (Figures 6G and 6H), the stand-alone iLV-to-BV conversion hardly occurred after brain vascular damages (Figures 6I-6K). Consequently, early BV regeneration failed to occur at 4 dpt and the majority of injured heterozygous animals died at 6 dpt (Figures 6L-6N). To confirm the significance of iLV-to-BV conversion for the survival of injured animals, two additional approaches were carried out to inhibit the transdifferentiation of stand-alone iLVs. Both LEC-specific overexpression of dnMAML-

Flag (Figures 6O-6Q) and general overexpression of EphrinB2a using the *hsp70l:ephrinB2a-p2A-Venus* transgene (Figures 6R-6T) led to defects in early BV regeneration and became lethal at 6 dpt. Taken together, these results demonstrate that early recovery of basic blood flows achieved by the stand-alone iLV-to-BV transdifferentiation is necessary for survival of the injured animals, which is the prerequisite to fulfill the late wave of BV regeneration.

### The iLV-to-BV Conversion Occurs in Adult Photochemical Thrombosis Model

The studies above were carried out using the zebrafish NTR-Mtz injury model at the larval stages. To answer the question whether the stand-alone iLV-to-BV transdifferentiation occurs in adults or in other brain vascular injury models, a pilot study in adults using photochemical thrombosis (Yu and Li, 2016) to mimic human cerebral thrombosis was performed (Figure 7A). At one day post injection of Rose bengal and cold light illumination (dpi), damages to regional brain vasculature in the illuminated areas, hereafter focusing on the cerebellum area, could be efficiently induced (Figures 7B and 7C). One day later at 2 dpi, ingrowth of BLECs into the injured area to form iLVs was observed, whereas BLECs in the uninjured brain exhibited interspersed distribution on the brain surface (Figures 7D-7G). At 3 dpi, the BEC-specific Dendra2 became detectable in a portion of *lyve1b*+ iLVs (Figures 7H-7L), indicating transdifferentiation of these iLVs to BVs. To validate this transdifferentiation in adults, the *Tg(kdrl:DenNTR; lyve1b:CreER^T2^; β-actin2:loxP-STOP-loxP-GFP)* lineage tracing line were subjected to photochemical thrombosis and 4-OHT administration. At 7 dpi, in contrast to the uninjured brain in which only BLECs were positive for GFP, approximately 15% of the Dendra2+ BVs were positive for GFP in the injured brain (Figures 7M-7P), indicating their origination from the *lyve1b*+ BLECs. The ratios of BVs derived from BLECs in adults (Figure 7P) were half of those in larvae (Figure 1D), which might be resulted from more survived remaining BVs in adults than in larvae after injury. This pilot study in adults suggests that the iLV-to-BV conversion is a conserved phenomenon in the adult photochemical thrombosis model.

**Figure 7.**
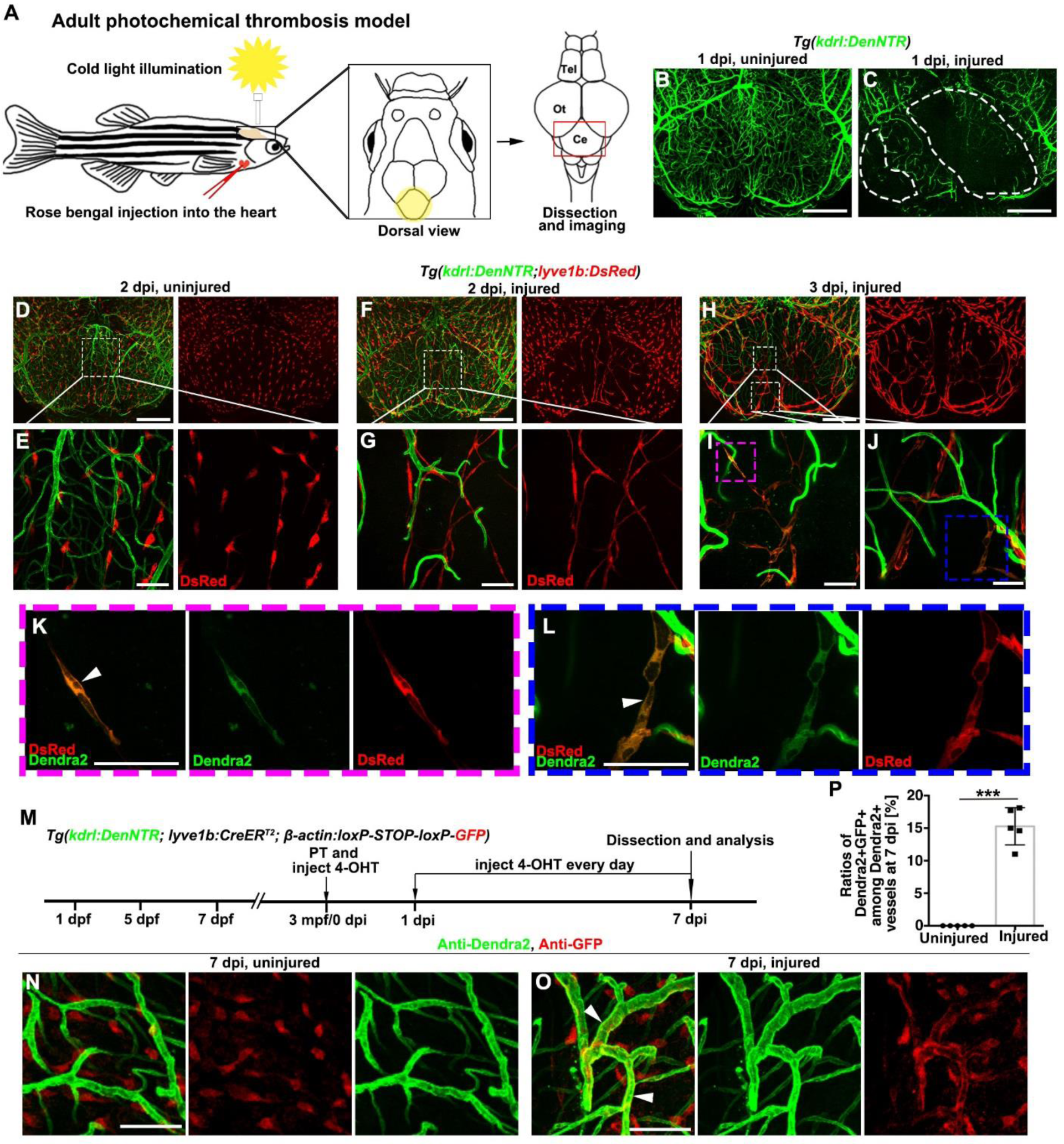
The iLV-to-BV conversion is conserved in the adult photochemical thrombosis model. (A) Illustrations of the cold light-induced thrombosis in the zebrafish brain. The yellow circle and red frame respectively indicate the illuminated and imaged cerebellum areas. Tel, Telencephalon; Ot, Optic tectum; Ce, Cerebellum. (B and C) In contrast to the uninjured control (B), damages to the cerebellar blood vasculature was observed at 1 day post illumination (dpi) (C). The dash lines mark the injured areas. n=10 adult brains. (D-G) At 2 dpi, in contrast to the interspersed distribution of BLECs on the surface of uninjured brain (D, E), ingrowth of BLECs into the injured area to form iLVs was observed (F, G). The framed areas in (D) and (F) are enlarged in (E) and (G), respectively. n=10 adult brains. (H-L) At 3 dpi, a portion of *lyve1b*+ iLVs in the injured cerebellum co-expressed the BEC-specific Dendra2 (K, L, arrowheads). From H to I and J, then to K and L, sequential enlargements of the framed areas are shown. n=10 adult brains. (M-P) Transgenic lines, time points of photochemical thrombosis (PT) and 4-OHT administration were shown in m. In contrast to the uninjured adult brain (N, injected rose bengal without cold light illumination, then injected 4-OHT from 0 dpi to 7 dpi every day), a portion of BVs were double positive for anti-GFP and anti-Dendra2 in the injured brain at 7 dpi (O, arrowheads). The statistics show the ratios of Dendra2+GFP+ vessels among all the Dendra2+ blood vessels at 7 dpi (P, n=5 regions from 5 different adult brains, Two-tailed unpaired *t*-test, ***, *p*<0.0001). Scale bar in B, C, D, F, H, 200 μm; Scale bar in E, G, I-L, N, O, 50 μm. Data are represented as mean±SD

## DISCUSSION

After cerebrovascular injury, the iLVs that rapidly ingrow into the injured brain parenchyma can be subdivided into two subpopulations, the stand-alone iLVs and track iLVs. The former is directly transdifferentiated into early-regenerated BVs to provide the ischemic tissues basic blood flows, thus enabling animal survival and fulfillment of late regenerations. The latter maintains its lymphatic fate and acts as “growing tracks” for late-regenerated BVs. The iLV transdifferentiation is dependent on the activation of Notch signaling, which is inhibited by the active EphB4a in the track iLECs.

EphrinB2a/EphB4a and the downstream Notch function as key regulatory molecules of lymphatic transdifferentiation vs maintenance (Figure S7).

In response to intensive brain vascular injury, iLVs keep a smart balance among different roles. One portion of iLVs are converted to early-regenerated BVs to provide prompt basic blood flows, while the other portion keep carrying out the lymphatic function of fluid drainage to continuously resolve the injury-caused brain edema. Both functions are indispensable to keep the injured animals survive, further enabling fulfillment of late BV regeneration to rebuild the brain vasculature.

Here, the formation of early-regenerated BVs is achieved via lymphangiogenesis plus transdifferentiation rather than via angiogenesis, indicating that angiogenesis is not the only route to form BVs. Lymphangiogenesis plus transdifferentiation can serve as an alternative. It is possible that the same regulatory molecule or signaling pathway plays distinct or even opposite roles in these two routes of BV formation. For example, *dll4* is expressed in BECs and the BEC Notch signaling limits angiogenesis during development (Bussmann et al., 2011; Hasan et al., 2017). Instead of this negative role in the developmental angiogenesis, Notch signaling plays positive role in the LV-to-BV transdifferentiation and thereby promote the prompt formation of early-regenerated brain BVs. The membrane ligand/receptor Delta/Notch could paracellularly activate the downstream signaling between two neighboring endothelial cells (Villa et al., 2001; Torres-Vazquez et al., 2003). After injury, expressions of both the ligand Dll4 and the receptor Notch1a as well as the Notch functional reporter and downstream target gene in the stand-alone, transdifferentiating iLVs implicate that the activation of Notch signaling in the stand-alone iLVs acts in a paracellular manner between two neighboring stand-alone iLECs.

LECs including BLECs originate from the BEC-to-LEC transdifferentiation during embryonic development (Srinivasan et al., 2007; Sabin, 1902; Schulte-Merker et al., 2011; Yaniv et al., 2006; Nicenboim et al., 2015; Venero Galanternik et al., 2017). But the reverse transdifferentiation, LEC-to-BEC, has hitherto been unappreciated for any biological function. In addition to shedding lights on the cellular and molecular mechanisms underlying prompt recovery of basic blood flows, this study also highlights the occurrence and biological significance of the LEC-to-BEC transdifferentiation in living vertebrates. Zebrafish iLVs come from BLECs (Chen et al., 2019), possibly corresponding to the mammalian LLECs (Shibata-Germanos et al., 2020). Our results suggest LLECs, the lymphatic transdifferentiation process, and the key regulatory molecules including EphrinB2a/EphB4a/Notch as potential post-ischemic therapeutic targets for enhancement of prompt brain vascular regeneration.

## ACKNOWLEDGMENTS

We thank PS Crosier, ND Lawson, and N Mochizuki for plasmids and fish lines. This work was supported by the National Natural Science Foundation of China 31730060, 91739304, and 32000576), the Natural Science Foundation of Chongqing (cstc2020jcyj-msxmX0882), and the 111 Program (B14037) (to L.L.).

## AUTHOR CONTRIBUTIONS

L.L., J.C. and J.H. designed the experimental strategy, analyzed data, and wrote the manuscript. X.L. performed the ephrin-related experiments. R.N. performed CLEM. Q.C. generated the *Tg(hsp70l:NICD-HA)* line. Q.Y. helped analyze data. J.C. performed all the other experiments in the study.

## STAR★METHODS

### LEAD CONTACT AND MATERIALS AVAILABILTY

#### Materials Availability Statements

Further information and requests for reagents may be directed to and will be fulfilled by the Lead Contact, Lingfei Luo. All zebrafish lines and plasmids generated in this study will be made available on reasonable request, but we may require a payment or a completed Material Transfer Agreement if there is potential for commercial application.

### EXPERIMENTAL MODEL AND SUBJECT DETAILS

#### Zebrafish husbandry and strains

Zebrafish strains were raised and maintained under standard laboratory conditions according to Institutional Animal Care and Use Committee protocols. Embryos were treated with 0.003% 1-phenyl-2-thiourea (PTU, Sigma) to inhibit pigment formation. All experimental protocols were approved by the Southwest University (Chongqing, China). The zebrafish facility and studies were approved by the Institutional Review Board of Southwest University (Chongqing, China). All results involved in the study were acquired according to ethical standards.

In NTR/Mtz models, the sex of zebrafish aged from 3 dpf to 12 dpf was unknown. In adult photochemical thrombosis model, both male or female fish were used for this experiment. A complete list of the zebrafish strains is provided in the Key Resources Table. All the strains are used as stable, germline transgenic lines in this study.

## METHOD DETAILS

### Molecular cloning and generation of transgenic lines

To generate the *pBluescript-kdrl:loxP-STOP-loxP-H2b-GFP-cryaa-CFP* construct, the *loxP -STOP-loxP-H2b-GFP-cryaa-CFP* fragment was subcloned downstream of *kdrl* promoter. The NICD sequence containing an HA tag was amplified from cDNA using the primers 5’-ATGTCCAGGAAGAGGAAGCGGGAAC-3’ and 5’-TTAAGCGTAATCTGGAACATCGTATGGGTACTTGAAGGCCTCTGGAATATG-3’, then subcloned into the *hsp70l:cryaa-CFP* vector to construct the *pBluescript-hsp70l:NICD-HA-cryaa-CFP*. To construct the plasmid *hsp70l:loxP-STOP-loxP-NICD-HA-cryaa-CFP*, the *H2b-GFP* fragment on the *hsp70l:loxP-STOP-loxP-H2b-GFP-cryaa-CFP* vector was replaced by the *NICD-HA* fragment. The *dnMAML* sequence containing a Flag tag was amplified from the genomic DNA of the *Tg(hsp70l:dnMAML-GFP)* embryos using the primers 5’-ACCGGTTATAGGGCTCGAGGCCACCATG-3’ and 5’-CTACTTGTCGTCATCGTCTTTGTAGTCCTGCCTGTGTTTGTTGGAGCGC-3’. The PCR product was cloned into the *pGEMT-easy* vector, followed by replacement of the *NICD-HA* fragment by the *dnMAML-Flag* fragment to construct the *hsp70l:loxP-STOP-loxP-dnMAML-Flag-cryaa-CFP* plasmid. For construction of the *pBluescript-hsp70l:ephrinB2a-P2A-Venus-cryaa-CFP* plasmid, the full-length CDS of zebrafish *ephrinB2a* was amplified from cDNA using the primers 5’-ACCGGTATGGGCGACTCTTTGTGGAGA-3’ and 5’-TCGCTCACTGACTCGCTGCGCT-3’, then ligated to the *p2A-Venus* sequence by overlap PCR and subcloned into the *hsp70l:cryaa-CFP* vector.

For constructing the *hsp:70l:loxP-STOP-loxP-EphB4aEE-p2A-Venus* plasmid, we first amplified the full-length zebrafish *ephB4a* cDNA using the primers 5’-CCCGGGATGGAGCTCTTCTCCAGGAA-3’ and 5’-GTACAGCACATTCCCTGGTGC-3’.

Then the *ephB4a* sequence overlapped with the *p2A-Venus* sequence was cloned into pGEMT vector. The *ephB4aEE* (Y596E, Y602E) was generated by the site-directed mutagenesis (Zheng et al., 2004). Then, the *ephB4aEE-p2A-Venus* were inserted into the *hsp70l:loxP-STOP-loxP;cryaa-cerulean* vector between the *Age*Ⅰ and *Spe*Ⅰ sites to construct the *pBluescript-hsp70l:loxP-STOP-loxP-EphB4aEE-p2A-Venus-cryaa-cerulean*. The successful mutated plasmids were identified by sequencing.

The *-5.2lyve1b:GFP* plasmid (Okuda et al., 2012) was co-injected with the capped *Tol2* transposase RNA (40-50 pg) for the transgenesis of *Tg(lyve1b:GFP)^cq86^*. The *Tg(kdrl:loxP -STOP-loxP-H2b-GFP;cryaa:CFP)^cq87^* was generated using the construct flanked by the *I-Sce*I restriction sites, which was co-injected with *I-Sce*I (NEB) into the *Tg(kdrl:CreER^T2^)^cq24^* embryos for transgenesis and screen. The constructs *pBluescript-hsp70l:loxP-STOP-loxP-NICD-HA;cryaa-CFP*, *pBluescript-hsp70l:loxP-STOP-loxP-dnMAML-Flag; cryaa-CFP*, *pBluescript-hsp70l:NICD-HA; cryaa-CFP*, *pBluescript-hsp70l:ephrinB2a-P2A-Venus; cryaa-CFP* ; and *pBluescript-hsp70l:loxp-STOP-loxp-EphB4aEE-p2A-Venus-cryaa-cerulean* were co-injected with *I-Sce*I (NEB) into the one-cell stage of embryos under the AB genetic background for transgenesis. For the maintenance of transgenic lines, animals were outcrossed at least every other generation to ensure genetic diversity.

### Generation of zebrafish genetic mutants and genotyping

The CRISPR-Cas9 technique was applied to generate the *ephrinB2a^cq101^* mutant. The target sequence of *ephrinB2a* gRNA 5’-CTTTGTGGAGAATATTACTT<u>TGG</u>-3’ (PAM site underlined) was located in the exon 1. Zebrafish zCas9 mRNA and the *ephrinB2a* gRNA were synthesized as described (Chang et al., 2013). After co-injection of zCas9 mRNA (300 pg) and *ephrinB2a* gRNA (50 pg) into 1-cell stage embryos, the genomic region flanking gRNA target site was amplified with a pair of gene specific primers (Key Resources Table) and sequenced for validation. The validated embryos were raised to adults (F0). The F0 fishes were screened to identify the founders whose progenies carried indels in the *ephrinB2a* gene. Then, the offspring of the identified founders was raised up. Individual F1 adults were genotyped by sequencing.

For genotyping the *ephB4a^hu3445^* mutants (Choe and Crump, 2015), the *dll4^j16e1^* hets (Leslie et al., 2007), and the *ephrinB2a^cq101^* mutants, genomic DNAs were amplified with primers (Key Resources Table). The amplified DNA fragments were subjected to high-resolution gel electrophoresis and sequencing.

### Antibody staining, combination of FISH and antibody staining

Whole-mount antibody staining and combination of FISH and antibody staining were performed as previously described (Chen et al., 2019; Liu et al., 2016; He et al., 2020) using antibodies against DsRed (1:200, Santa-Cruz), GFP or Venus (1:500, Abcam and Santa Cruz), Dendra2 (1:500, Antibody-online), Prox1 (1:1000, Abcam), and phospho-EphB4a (1:250, Signalway Antibody). Secondary antibodies used in the study included donkey anti-goat IgG Alexa fluor 488-conjugated (1:1000, Invitrogen#A11055), donkey anti-goat IgG Alexa fluor 405-conjugated (1:1000, Abcam#ab175664), donkey anti-mouse IgG Alexa fluor 568-conjugated (1:1000, Invitrogen#A10037), donkey anti-rabbit IgG Alexa fluor 647-conjugated (1:1000, Invitrogen#A31573), donkey anti-mouse IgG Alexa fluor 647-conjugated (1:1000, Invitrogen#A31571), donkey anti-rabbit IgG Alexa fluor 568-conjugated (1:1000, Invitrogen#A10042), donkey anti-rabbit IgG Alexa fluor 488-conjugated (1:1000, Invitrogen, # A21206), and goat anti-rabbit IgG Alexa fluor 405-conjugated (1:1000, Invitrogen#A31556).

### TUNEL assay

Larvae were fixed in 4% paraformaldehyde in PBS at 4 °C overnight, subjected to skin removal, and assayed using the *In Situ* Cell Death Detection Kit, TMR Red (Roche) according to the manufacturer’s instruction.

### Temporal control of CreER activities

4-hydroxytamoxifen (4-OHT, Sigma) was dissolved in 100% ethanol to prepare a stock concentration of 10 mM. Transgenic larvae were incubated in 5 μM working solution for 24 hours from before Mtz treatment stage (3 dpf) to 1 dpt (4 dpf), followed by three washes with fresh egg water.

### Chemical treatment

Mtz treatment was performed as previously described (Chen et al., 2019), 1 mM Mtz (Sigma) dissolved in 0.2% DMSO was used for treatment for 5 hours, after treatment, the larvae were washed three times with egg water, and recovered in the egg water with 0.003% PTU. Control larvae were incubated in 0.2% DMSO in egg water with PTU.

50 mM DAPT (Sigma) stock solution in DMSO was used to prepare a working solution. The larvae were incubated in 50 μM DAPT working solution at the indicated time frame (He et al., 2014). Control larvae were incubated in 0.1% DMSO in egg water with PTU.

### Heat shock and mosaic ectopic expressions

The *Tg(hsp70l:NICD-HA;cryaa:CFP)* and *Tg(hsp70l:dnMAML-GFP)* larvae were screened for the lense-expressing CFP for general heat shock experiments. Heat shock was performed at 38.5 °C for 40 minutes at the indicated time frame, followed by incubation at 28.5 °C for further raising.

For the LEC-specific overexpression of NICD-HA, dnMAML-Flag, and EphB4aEE-P2A-Venus, the *Tg(lyve1b:CreER^T2^; hsp70l:loxP-STOP-loxP-NICD-HA; cryaa-CFP)* or *Tg(lyve1b:CreER^T2^; hsp70l:loxP-STOP-loxP-dnMAML-Flag; cryaa-CFP)*, and *Tg(lyve1b:CreER^T2^; hsp70l:loxP-STOP-loxP-ephB4aEE-p2A-Venus; cryaa-CFP)* larvae were first screened for the lense-expressing CFP. After incubation with 4-OHT from 3 dpf to 1 dpt/4 dpf, heat shock was carried out at 38.5 °C for 40 minutes. Then, heat-shock was repeatedly performed every 24 hours from 1 dpt to 4 dpt. The larvae were fixed at the indicated stage for the following assays, all the larvae were identified by genotyping or antibody staining.

For ectopic expression of EphrinB2a, the *hsp70l-ephrinB2a-p2A-Venus* or *hsp70l-venus* plasmid and *I-Sce*I (NEB) were co-injected into the blastomeres at the one-cell stage of *Tg(kdrl:DenNTR; lyve1b:DsRed)*. Larvae at specific stages (from 2 dpt to 4 dpt) were heat-shocked at 37.5 °C for 45 minutes and return to 28.5 °C for further raising.

The larvae were fixed at the indicated stage for the antibody staining.

### Confocal imaging and CLEM

Antibody, FISH, and TUNEL stained larvae were mounted in 1.2% low melting point agarose and imaged using ZEN2010 software equipped on an LSM880-Airyscan confocal microscope (Carl Zeiss). For time-lapse live imaging, larvae were mounted in 1.2% low melting point agarose in the egg water with 0.003% PTU using 35-mm glass bottom dishes. Time-lapse images were captured using a 20× water immersion objective mounted on the LSM880-Airyscan confocal microscope equipped with a heating stage to maintain 28.5 °C. Z image stacks were collected every 30 minutes, and three-dimensional data sets were compiled using ZEN2010 software (Carl Zeiss). The CLEM was performed using an FIB-SEM Crossbeam 540 (Carl Zeiss) as previously described (Chen et al., 2019).

### Dye injection, vibratome section of adult brains and antibody staining

The dye injection was performed as previously described (Chen et al., 2019; van Lessen et al., 2017). In brief, a suspension of IgG-conjugated Alexa fluor 647 (2 mg/ml, A31573, Invitrogen) was intracerebroventricularly injected using glass capillary needles.

Adult zebrafish brains were dissected, fixed in 4% PFA at 4 °C overnight, and embedded in 4% low melting point agarose, followed by 50 μm sections using VT1000S vibratome (Leica). These sections were subjected to antibody staining using the anti-DsRed (1:1000, Santa-Cruz, sc101526) and anti-Dendra2 (1:1000, Antibody-online, #ABIN361314) primary antibodies. Then, the donkey anti-rabbit IgG Alexa fluor 488-conjugated (1:1000, Invitrogen, # A21206) and donkey anti-mouse IgG Alexa fluor 568-conjugated (1:1000, Abcam, # ab10037) secondary antibodies were used to label Dendra2 and DsRed, respectively.

### Photoconversion

For photoconversion, the *Tg(lyve1b:Kaede; kdrl:CFP-NTR)* larvae were mounted in 1.2% low melting point (L.M.P.) agarose. The focused BLECs loop regions of 1 dpt were irradiated for 30 seconds by the 405-nm laser. The Kaede epifluorescences were converted from green to red after photoconversion.

### Induction of adult photochemical thrombosis and whole-mount imaging

Adult fishes (older than 3 months post fertilization, males or females) were anesthetized with 140 mg/L tricaine (MS-222) and placed upside down into a sponge slit moistened with tricaine water. Rose bengal (8mg/ml, 632-69-9, Sangon Biotech) or 4-OHT (2mM) was injected into the heart with a glass capillary needle. The total volume of injection should be less than 40 nl. After injection, the fish was placed upright into another sponge slit moistened with 75 mg/L tricaine. The cerebellum region was exposed to cold light for 3-5 minutes as previously reported (Yu and Li, 2016). The illuminator and the light probe were purchased from Carl Zeiss (CL 6000 LED).

After illumination, brain of adult fish at the indicated stage were dissected and placed into cold PBS for imaging. The dissected brains were whole-mount imaged by an LSM880 confocal microscope (Carl Zeiss) equipped with 10X Air (0.45 N.A.) and 20X water immersion (0.95 N.A.) objectives using ZEN 2010 software. The large size of the adult zebrafish brains often required tile scan acquisitions that were later stitched using ZEN 2010 software.

## QUANTIFICATION AND STATISTICAL ANALYSIS

All statistical calculations were performed using GraphPad Prism. Variance for all groups of data are presented as ±S.D. Larvae were collected from incrosses and outcrosses of several pairs of adult zebrafish. The *ephB4a^hu3445^* mutant, *ephrinB2a^cq101^* heterozygotes and *dll4^j16e1/+^* heterozygotes were raised to adulthood. More than six pairs of fish were used to set one specific cross. For a cross between *ephB4a^hu3445^* homozygous parents, more than 30 larvae were used for analyses in each treatment. For a cross between *dll4^j16e1/+^* heterozygous and *ephrinB2a* heterozygotes parents, more than 50 larvae were used for each treatment and subjected for genotyping.

Genotyping preceded phenotyping in homozygotes and heterozygotes analyses. Hence, analyses were genotypy non-blinded. In the other experiments, the investigators were not blinded to group allocation during data collection and/or analysis. All experiments comparing treatment groups were carried out using randomly assigned siblings. After at least two repeated experiments, data were analysed for statistical significance using two-tailed unpaired *t*-test or Two-way ANOVA by Sidak’s and Dunnett’s multiple comparisons test. A value of *P*<0.05 was considered to be statistically significant. No data were excluded from analyses. The exact sample size (n), *P* value for each experimental group and statistical tests were indicated in the figure legends.

## Supplemental Video Legends

**Video S1. The stand-alone iLVs gradually express the BV marker and obtain blood flows at 3 dpt. Related to Figure 1** Living imaging of the *Tg(kdrl:DenNTR; prox1a^BAC^:KalTA4; UAS:TagRFP)* transgenic line at 2 dpt, 3 dpt, 4 dpt after Mtz treatment. The Alexa647-IgG was intracerebroventricularly injected before Mtz treatment and endocytosed by LECs. Note that the RFP+Alexa647+ iLVs (arrowheads) express Dendra2 and obtain blood flows at 3 dpt. Scale bar, 20 μm.

**Video S2. Time lapse live imaging for the real-time processes of iLV ingrowth and transdifferentiation. Related to Figure 1** At 24:00 (24 hpt), the DsRed+ iLV began to ingrow into the injured brain parenchyma. At 60:00 (60 hpt), the iLV began to express BEC-specific Dendra2, which became evident at 72:00 (72 hpt) and afterwards. The left and right panels of the video show DsRed/Dendra2 double channels and Dendra2 single channel, respectively. Duration of imaging was 96 hours. The elapsed time is indicated in hours:minutes post 0 dpt. Scale bar, 50 μm.

**Video S3. The stand-alone iLV derived from the photoconverted BLECs is converted to BV with blood flows. Related to Figure 1** Note that a portion of iLECs in the indicated iLV (arrowhead) originated from the photoconverted, Kaede-red BLECs. Blood flows could be observed. Scale bar, 20 μm.

**Video S4. The track iLVs are converted to BVs with blood flows in the *ephB4a* mutant. Related to Figure 3** In the *ephB4a* mutant under the *Tg(kdrl:DenNTR; lyve1b:DsRed)* transgenic background, the DsRed-positive track iLV expressed the BV-specific Dendra2 and finally obtained blood flows (arrowheads). The video above displays the bright field of the video below. Scale bar, 20 μm.

