## Supplemental Information for "Urgent Brain Vascular Regeneration Occurs via Lymphatic Transdifferentiation"

### Supplemental Figures

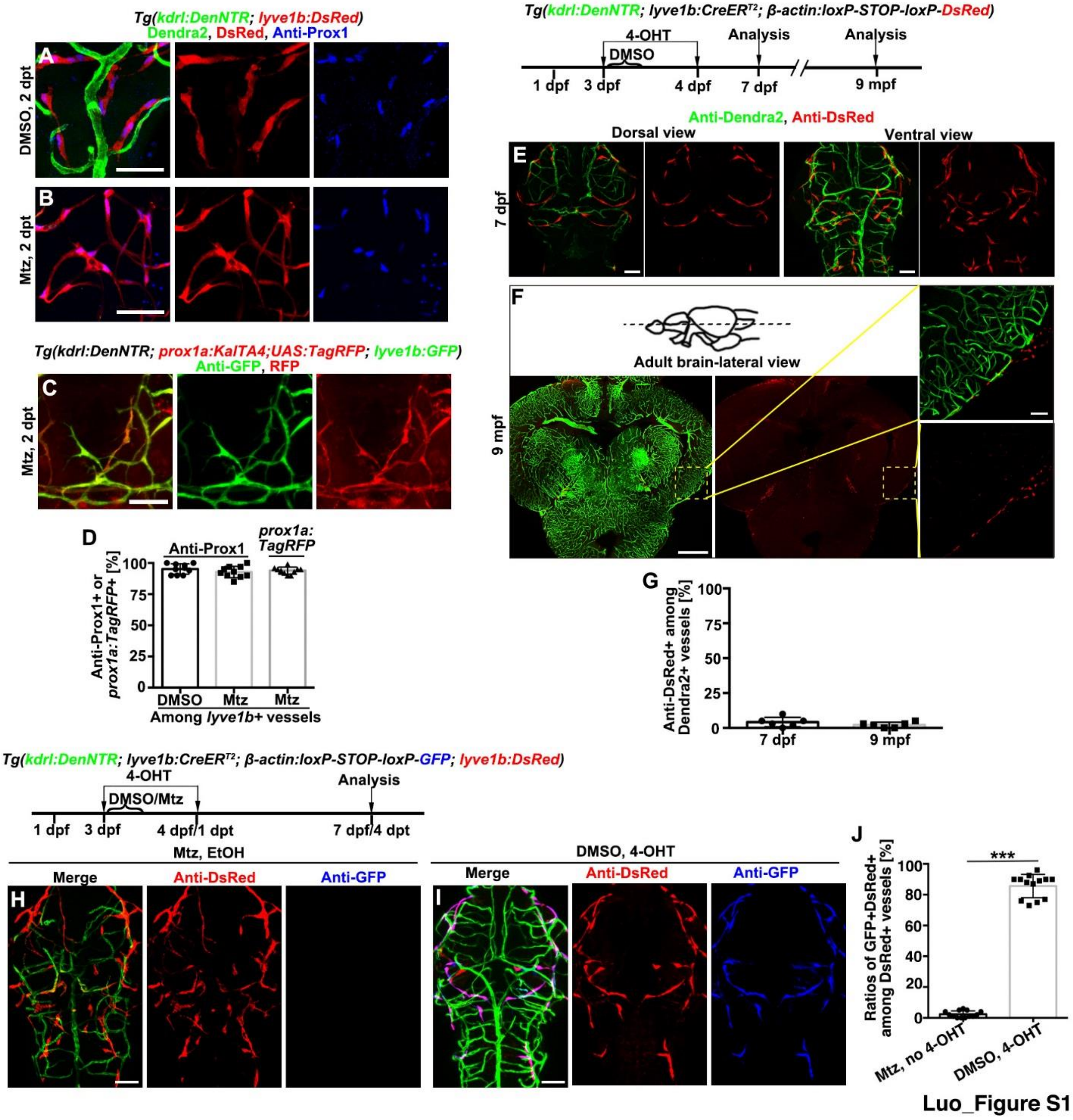


**Figure S1. Validations of the LEC-specific promoter inside the skull and the lineage tracing line, Related to Figure 1**

(A-D) In both control and injured larvae, all the *lyve1b:DsRed*-expressing BLECs (A) and iLVs (B) inside the skull were positive for anti-Prox1 antibody staining at 2 dpt. In the *Tg(kdrl:DenNTR; prox1a:KalTA4;UAS:TagRFP; lyve1b:GFP)* line, the GFP+ vessels completely overlapped with the TagRFP+ vessels at 2 dpt after Mtz treatment (C). The statistics show the ratios anti-Prox1+ or TagRFP+ among all the *lyve1b*+ vessels inside the skull (D, n=10 larvae).

(E-G) Treatment of the *Tg(kdrl:DenNTR; lyve1b:CreER*$^{T2}$*; β-actin2:loxP-STOP-loxP-DsRed)* line with 4-OHT from 3 dpf to 4 dpf labeled BLECs with DsRed. None of the Dendra2+ brain BVs were positive for anti-DsRed at 7 dpf (E) and 9 months post fertilization (mpt) (F). Sections of brain at 9 mpt are shown in (F). Scale bar, 400 μm. The framed areas are enlarged in right panels. Scale bar, 50 μm. The statistics show the ratios of vessels positive for anti-DsRed among all the Dendra2+ vessels in the brain of 7 dpf and 9 mpf (G, 7 dpf, n=6 larvae; 9mpf, n=6 sections from 6 different adult brains).

(H-J) The triple transgenic *Tg(lyve1b:CreER*$^{T2}$*; β-actin2:loxP-STOP-loxP-GFP; kdrl:DenNTR)* lineage tracing line was crossed with the *Tg(lyve1b:DsRed)* line, followed by treatment with Mtz plus ethanol as the negative control. No induction of GFP expression indicated no Mtz-caused CreER leakiness (H). The above crossed quadruple transgenic larvae were treated with DMSO plus 4-OHT as the positive control. More than 90% of the DsRed+ vessels were also positive for GFP, indicating high labeling efficiency of the LEC-derived cells by GFP (I). The statistics show the ratios of GFP+DsRed+ vessels among all the DsRed+ vessels (J, n=13 larvae, ***, $p<0.0001$).

Scale bar, 50 µm otherwise indicated. *p* values calculated using two-tailed unpaired *t*-tests.. Data are represented as mean±SD.

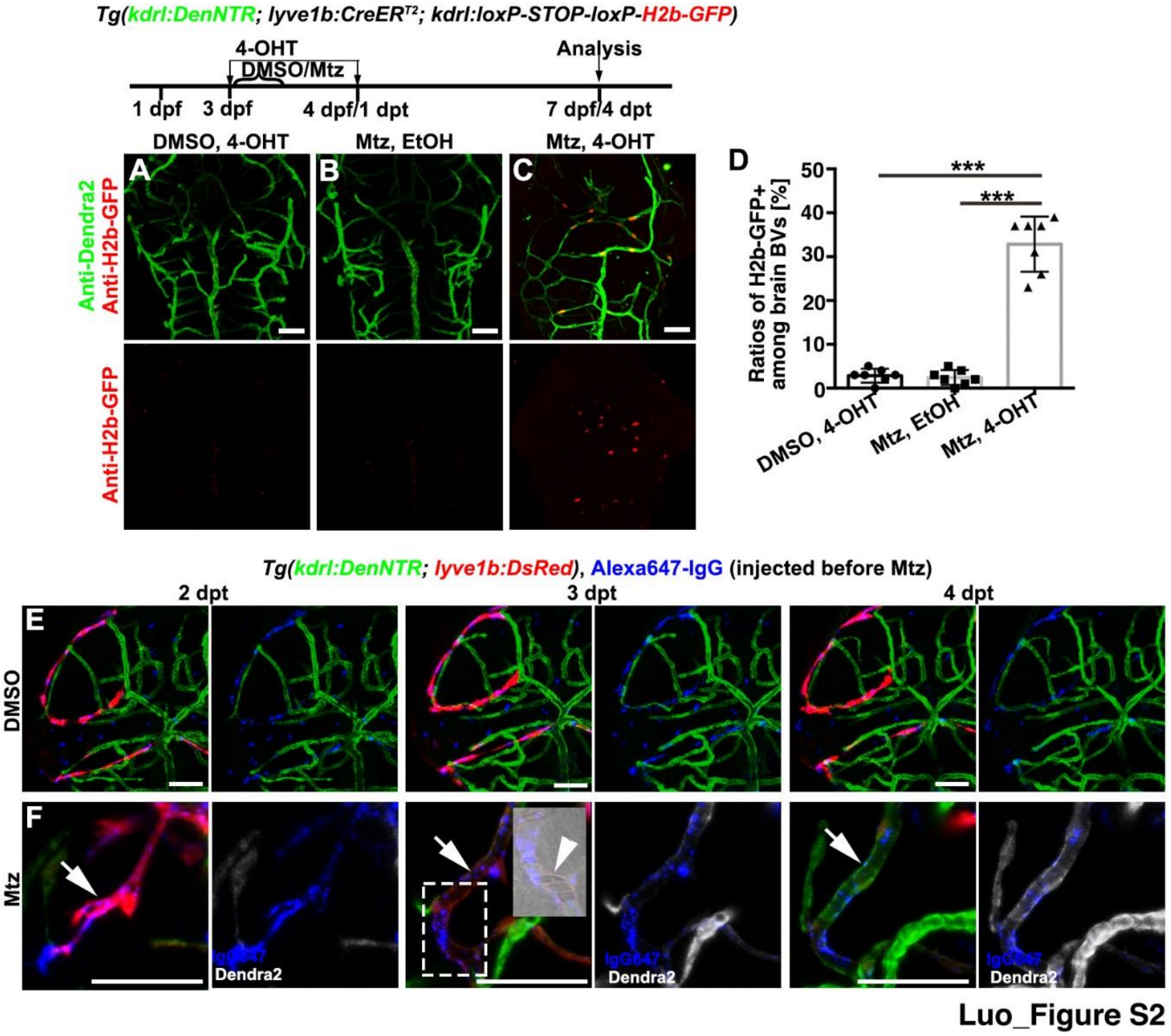


**Figure S2. Early-regenerated BVs come from iLV transdifferentiation, Related to Figure 1 and Figure 2.**

(A-D) Controlled by treatments with DMSO plus 4-OHT (A) and Mtz plus ethanol (B), treatment of the *Tg(kdrl:DenNTR; lyve1b:CreER$^{T2}$; kdrl:loxP-STOP-loxP-H2b-GFP)* line with Mtz plus 4-OHT showed that a portion of brain BVs at 4 dpt were positive for H2b-GFP (C). The statistics show the ratios of H2b-GFP+ vessels among Dendra2+ brain BVs (D, n=7 larvae, ***, $p$<0.0001).

(E and F) In contrast to the DsRed+Alexa647+ BLECs in the uninjured control (E), the

DsRed+Alexa647+ iLVs gradually expressed Dendra2 from 2 dpt to 4 dpt after Mtz treatment (F, arrows). The bright field image of the framed area is displayed and arrowhead indicates the recovered blood flows at 3 dpt. The Alexa647-IgG was injected before Mtz treatment.

Scale bar, 50 μm. $p$ values calculated using two-tailed unpaired $t$-tests.. Data are represented as mean±SD.

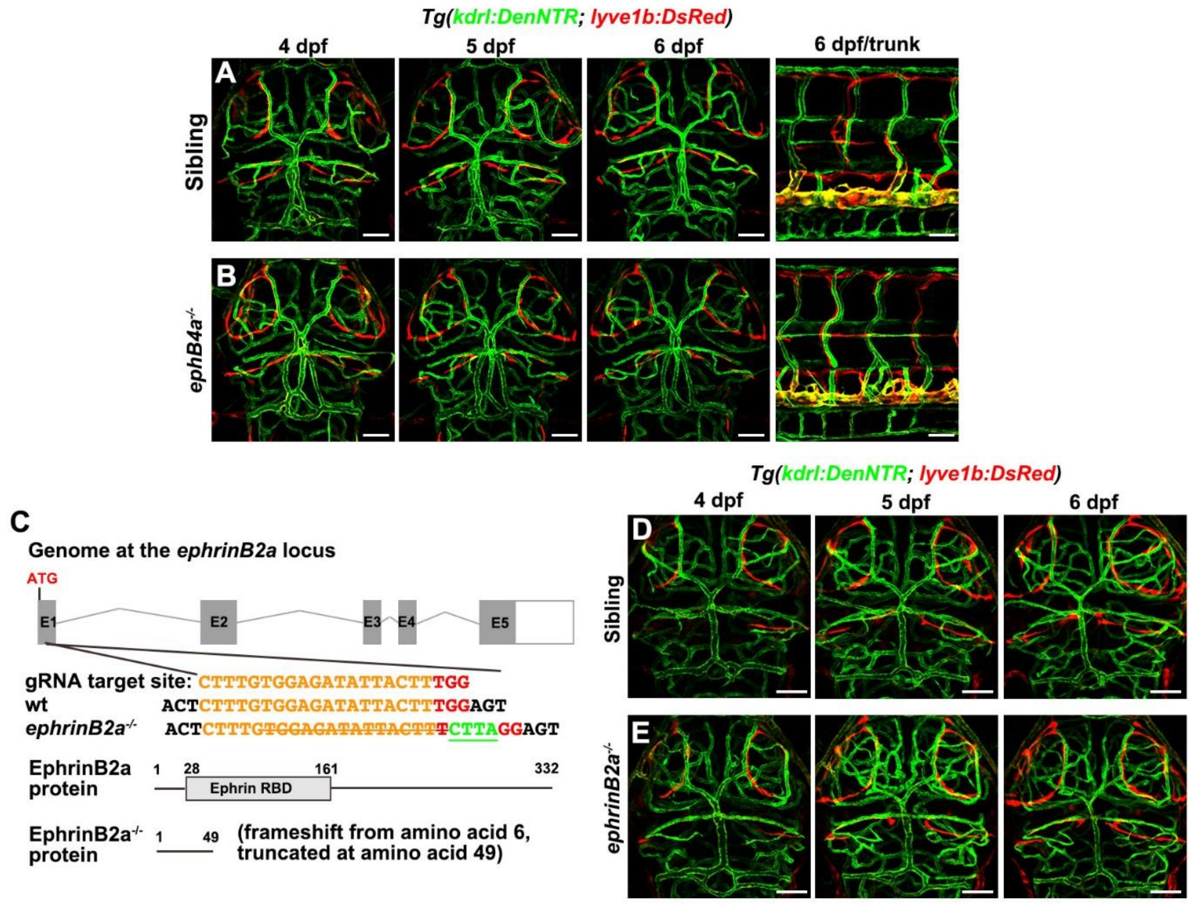


Luo_Figure S3

**Figure S3. The vascular development exhibits normal in *ephB4a* and *ephrinB2a* mutant. Related to Figure 3.**

(A and B) Compared with the siblings (A, n=20/20 larvae), the brain blood vascular and lymphatic development remained normal in the *ephB4a* mutant (B, n=25/25 larvae). (C-E) Schematic diagram showed generation of the *ephrinB2a* mutant. The orange letter, red letters, orange horizontal line, and underlined green letters denote the gRNA target site, the PAM site, the deletion of 15 bp, and insertion of 4 bp in the ephrinB2a gene, respectively. Note that the majority of the EphrinB2a receptor binding domain (RBD) becomes truncated in the mutant (C). In contrast to the siblings (D, n=30/30 larvae), the brain vascular and the BLECs development were normal in the *ephrinB2a* mutant (E, n=20/21 larvae). Scale bar, 50 μm.

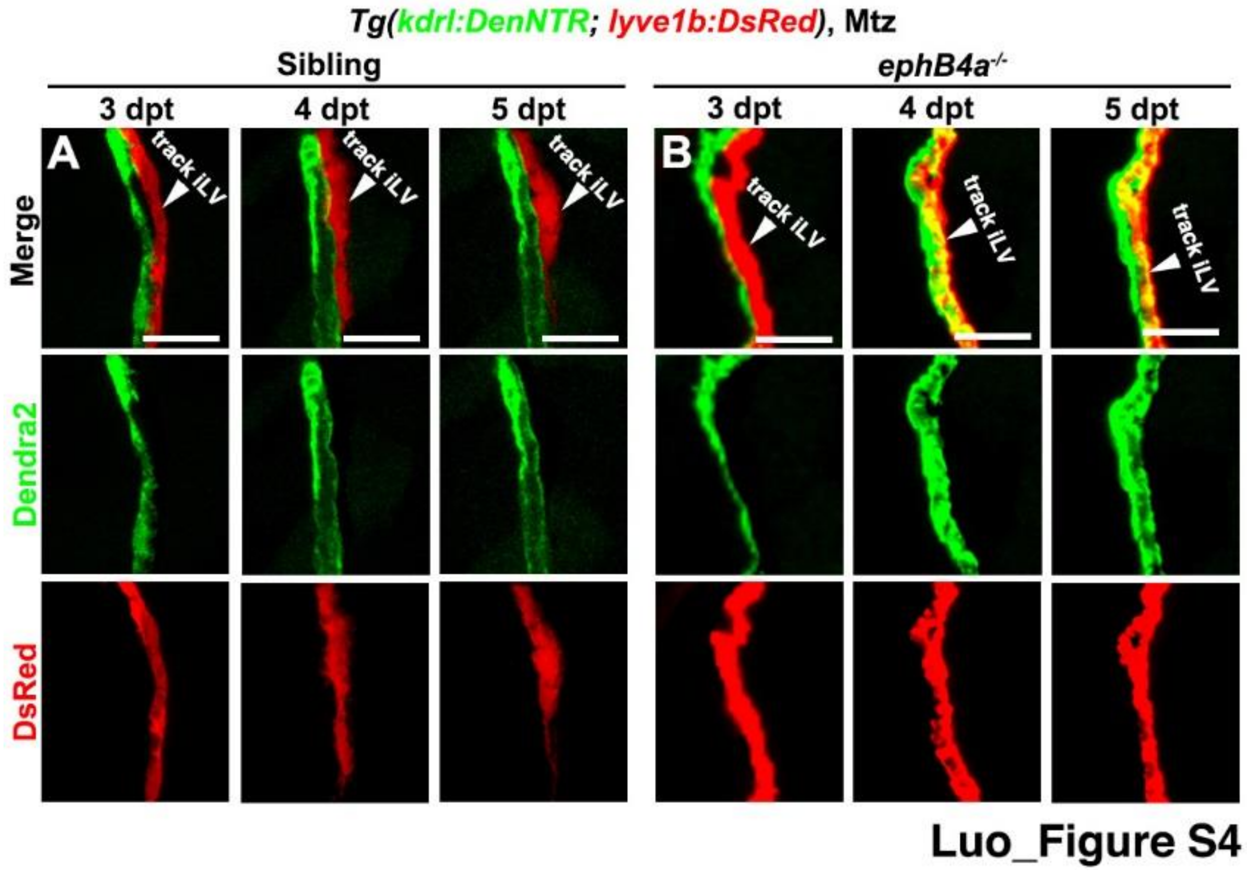


**Figure S4. Time-lapse imaging of the track iLV-to-BV conversion in *ephB4a* mutant. Related to Figure 3.**

(A and B) In contrast to the DsRed+Dendra2- track iLVs in the siblings (A), the DsRed+ track iLVs in the *ephB4a* mutant gradually expressed Dendra2 from 4 dpt (B). Scale bar, 20 μm.

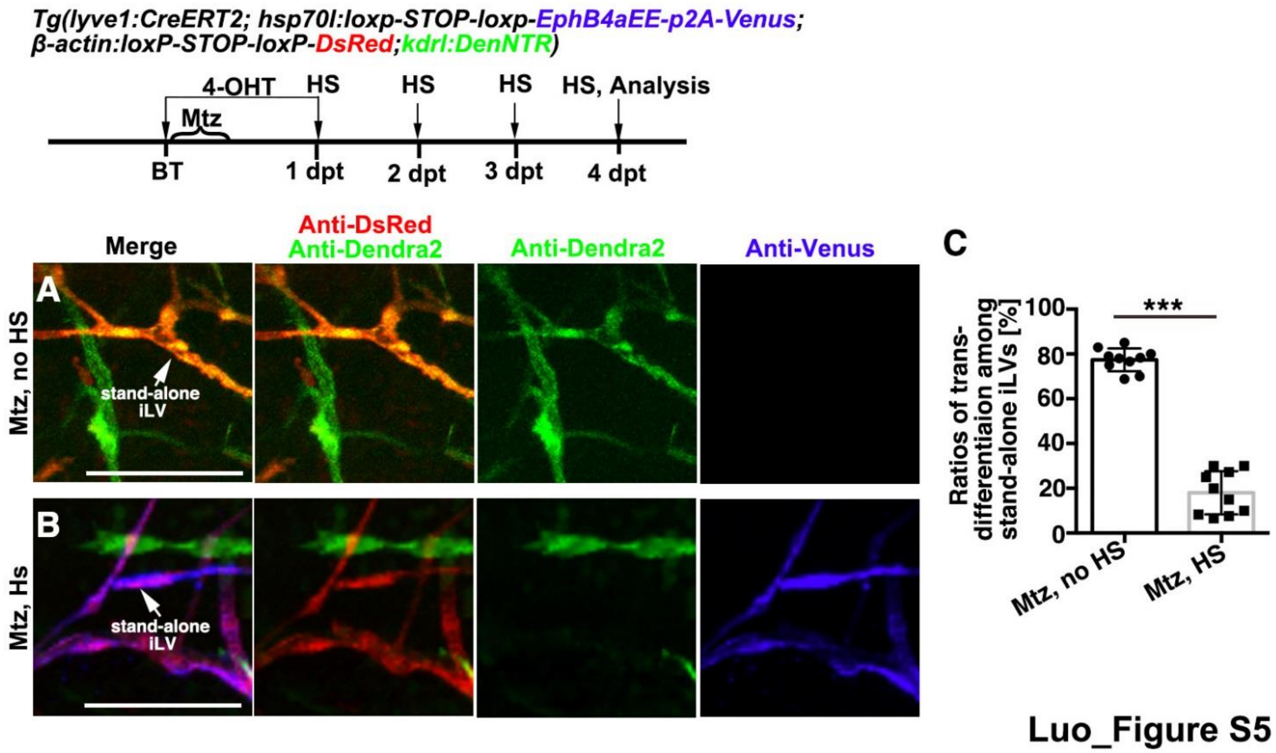


**Figure S5. The LEC-specific overexpression of EphB4aEE blocks the transdifferentiation of stand-alone iLV. Related to Figure 3.**

(A-C) In contrast to the transdifferentiating stand-alone iLVs (Dendra2+DsRed+) in the control (A), the heat shock-induced LEC-specific overexpression of EphB4aEE blocked the transdifferentiation of stand-alone iLVs (B). The statistics show the ratios of transdifferentiating stand-alone iLVs (Dendra2+DsRed+) among all the stand-alone DsRed+ iLVs (C, n=10 larvae, Two-tailed unpaired *t*-test. ***, $p<0.0001$). Scale bar, 50 μm. Data are represented as mean±SD.

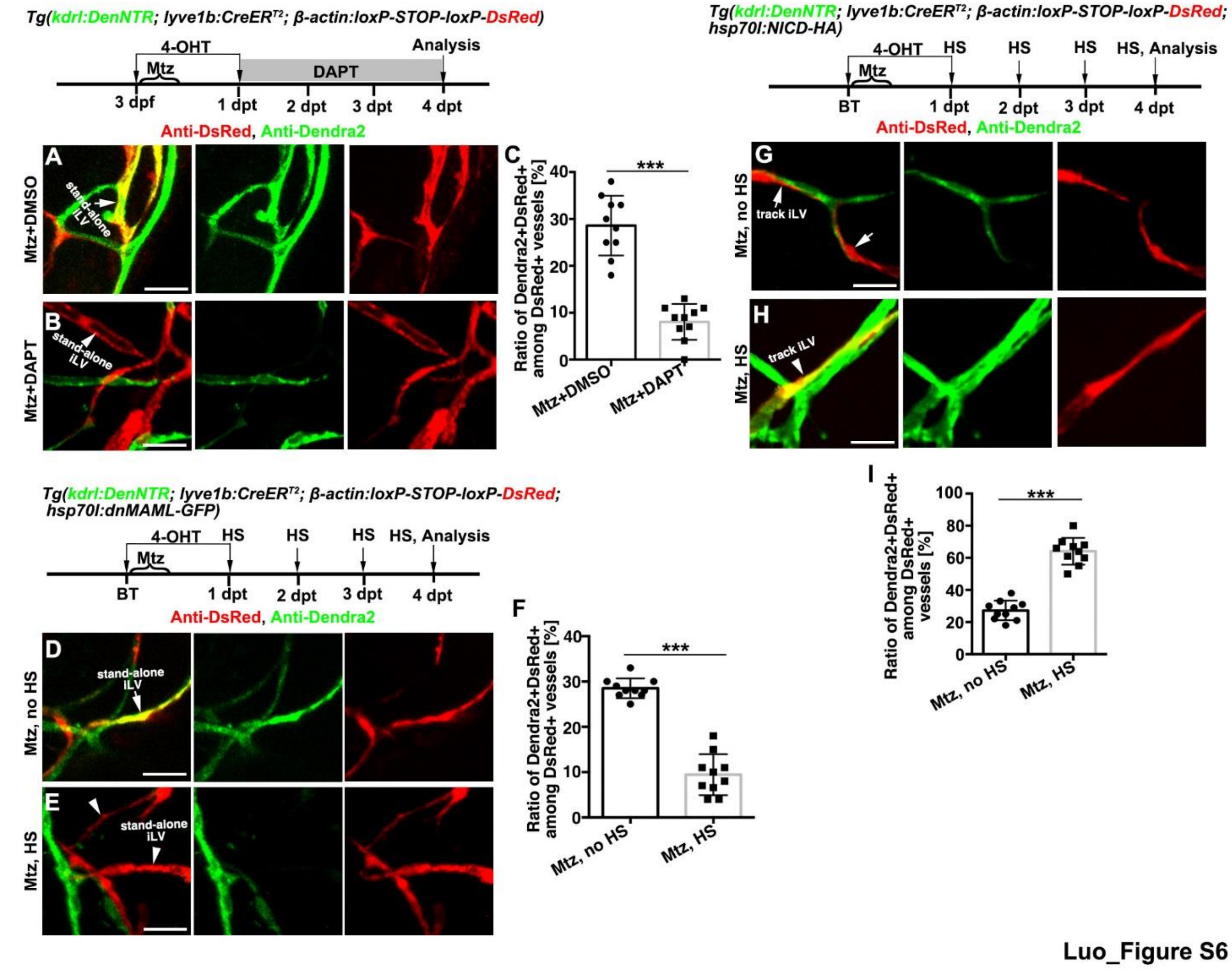


**Figure S6. Activation of Notch is required and sufficient to induce the iLV-to-BV transdifferentiation, Related to Figures 5 and 6**

(A-F) Transdifferentiation of stand-alone iLVs (A, D, arrows) was inhibited by the suppression of Notch signaling using DAPT (B, arrowhead) or overexpression of dnMAML-Flag (E, arrowheads). The statistics show the ratios of Dendra2+DsRed+ vessels among all the DsRed+ vessels (C, F, n=10 larvae, ***, $p$<0.0001).

(G-I) The track iLVs (G, arrows) were induced to undergo LV-to-BV transdifferentiation by the overexpression of NICD-HA (H, arrowhead). The statistics show the ratios of

Dendra2+DsRed+ vessels among all the DsRed+ vessels (**I**, n=10 larvae, ***, $p<0.0001$).

Scale bar, 20 μm. *p* values calculated using two-tailed unpaired *t*-tests.. Data are represented as mean±SD.

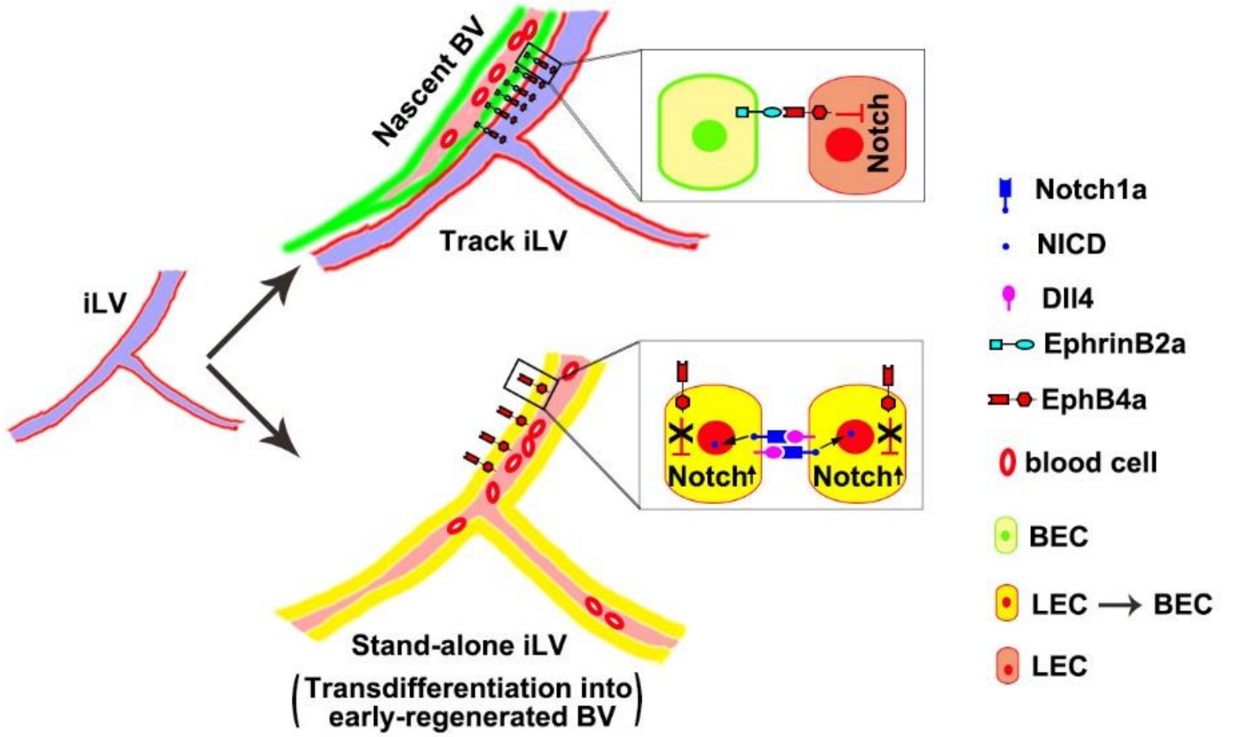


Luo_Figure S7

**Figure S7. Formation of early-regenerated BVs is achieved by transdifferentiation of stand-alone iLVs, Related to Figures 3 to 6**

Schematic illustrations of the iLV-to-BV transdifferentiation. After cerebrovascular injury, the iLVs that rapidly ingrow into the injured brain parenchyma can be subdivided into two subpopulations, the track iLVs and stand-alone iLVs. The track iLVs express EphB4a, which is paracellularly activated by the nascent BV-expressing EphrinB2a to suppress the activation of Notch in the track iLECs. By contrast, EphB4a is inactive in the stand-alone iLVs because no EphrinB2a ligand is available. So, the Notch signaling is derepressed in the stand-alone iLECs to induce LV-to-BV transdifferentiation, through which formation of early-regenerated brain BVs is achieved.
